# Different conformations of EGF-induced receptor dimers involved in signaling and internalization

**DOI:** 10.1101/2022.04.19.488777

**Authors:** Jordi Haubrich, Jurriaan M. Zwier, Fabienne Charrier-Savournin, Laurent Prézeau, Jean-Philippe Pin

**Author notes:** Author contributions: J.H., L.P. and J.-P.P. designed research and wrote the paper; F.C.-S. and J.M.Z. designed research; J.H. performed research and analyzed data.

## Abstract

The structural basis of the activation of EGF receptors (EGFR) is still a matter of debate despite the importance of this target in cancer treatment. Whether agonists induce dimer formation or act on pre-formed dimers remain discussed. Here we provide direct evidence that EGFR activation results from EGF-induced dimer formation. This is well illustrated by i) a large increase in time resolved (TR)-FRET between snap-tagged EGFR subunits induced by agonists, ii) a similar effect of Erlotinib-related TK inhibitors despite the inactive state of the binding domain of the subunits, and iii) a similar TR-FRET efficacy in EGFR dimers stabilized by EGF or erlotinib with binding domains in active and inactive states, respectively. Surprisingly, TK inhibitors do not inhibit EGF-induced EGFR internalization despite their ability to fully block EGFR signaling. Only Erlotinib-related TK inhibitors promoting asymmetric dimers could slow down this process, while the lapatinib-related ones have almost no effect. These results reveal that the conformation of the intracellular TK dimer, rather than the known EGFR signaling is critical for EGFR internalization. These results also illustrate clear differences in the mode of action of TK inhibitors on the EGFR.

---

In the group of receptor tyrosine kinases (RTKs) the Epidermal Growth Factor Receptor (EGFR) is one of the most-studied receptors. The EGFR is expressed in many tissues and plays a vital role in biological processes as apoptosis, cell growth and differentiation, migration and more ^1^.

*EGFR* gene mutations or amplification are found in many cancers, among which non-small cell lung cancer (NSCLC) has one of the lowest survival rates ^2^. Depending on histological and genotypical analysis of NSCLC, different and combinatorial treatment strategies are being used. The treatment strategy for patients with sensitizing EGFR mutations like a substitution of leucine for arginine at position 858 (L858R) involves ATP-competitive reversible first-generation TK inhibitors (e.g. erlotinib) ^3^. Resistance mechanisms occurring after treatment with first-generation TK inhibitors are common. In around 50% of these patients the apparition of a secondary mutation in the *EGFR* gene is prevalent (e.g. the substitution of a threonine for a methionine at position 790 (T790M)) ^4^. This mutation desensitizes first-generation TK inhibitors and induces higher enzymatic activity of the EGFR ^5^. Treatment options for L858R and T790M double-mutated EGFR (EGFR_NSCLC_) involves irreversible second and third-generation TK inhibitors (i.e. dacomitinib and osimertinib, respectively) ^6, 7^.

The EGFR has seven endogenous agonists, that are all small proteins ^8^. Upon binding to the extracellular domain, EGFR activators promote allosteric processes between two assembled EGFRs leading to the association of the intracellular tyrosine kinase (TK) domains, and the activation of one of them. EGFR activity is exclusively occurring when the two TK domains adopt an asymmetric conformation, i.e. there is an enzymatically inactive (Activator) and active (Receiver) domain ^9^. The Receiver domain initiates ATP-dependent phosphorylation of tyrosine residues on the intracellular domain of the Activator. Phosphorylation of its carboxy-tail residues leads to downstream signaling events such as activation of ERK1/2 and PI3K/AKT/mTOR ^10^. Conversely, receptor trafficking may be both dependent ^11^ and independent of the phosphorylation of carboxy terminal-tail residues ^12, 13^. Surprisingly, whether this process results from ligand-induced dimerization ^14^ or from conformational changes within pre-formed dimers ^15^ is still a matter of debate.

We have used techniques based on time-resolved (TR) Förster resonance energy transfer (TR-FRET) to analyze the mode of action of EGFR agonists and various types of TK inhibitors. We provide clear evidence that EGFR agonists promote dimer formation through direct contact between the extracellular domains, resulting in receptor phosphorylation and internalization. We show that first-generation TK inhibitors and dacomitinib also induce EGFR dimer formation, through a direct association of the TK domains, independently of the conformation of the extracellular domain. Surprisingly, TK inhibitors do not inhibit EGF-induced EGFR internalization, demonstrating EGFR signaling is not required for this process. Importantly, TK inhibitors inducing dimers slow down EGF-induced internalization of the EGFR, revealing a link between TK inhibitor-induced EGFR conformation and EGFR trafficking. These data reveal that differential effects induced by TK inhibitors can result from different conformation of the ligand-induced EGFR dimers.

## Results

### Intersubunit FRET induced by agonists and group I TK inhibitors through the TK domain

Intersubunit FRET was measured after randomly labelling the SNAP-tags with SNAP-Lumi4-Tb (donor) and SNAP-Green (acceptor) (Fig. 1A) ^16^ in monoclonal stable cell lines expressing SNAP-EGFR or SNAP-EGFR_NSCLC_. While a very low FRET was measured under basal condition (Fig. 1B, insert), agonists and the group I TK inhibitors, dacomitinib, erlotinib and PD153035 induced a large increase in intersubunit FRET (Fig. 1B, C and Suppl. Fig. 2 and 3A). Since FRET efficacy is related to distance of the FRET-pair, total FRET levels may change depending on the conformational changes and on the number of molecules in FRET. Group I TK inhibitors induced a lower TR-FRET signal than agonists, raising the question whether this is due to a different conformation or to a lower number of induced dimers. In contrast, the other TK inhibitors, GW583340 and lapatinib (group II inhibitors), osimertinib, cetuximab (i.e. an inhibitory antibody) and AG1024 (i.e. an insulin-like growth factor receptor inhibitor) did not induce intersubunit FRET (Fig. 1C and Suppl. Fig. 2 and 3A).

**Fig. 1.**
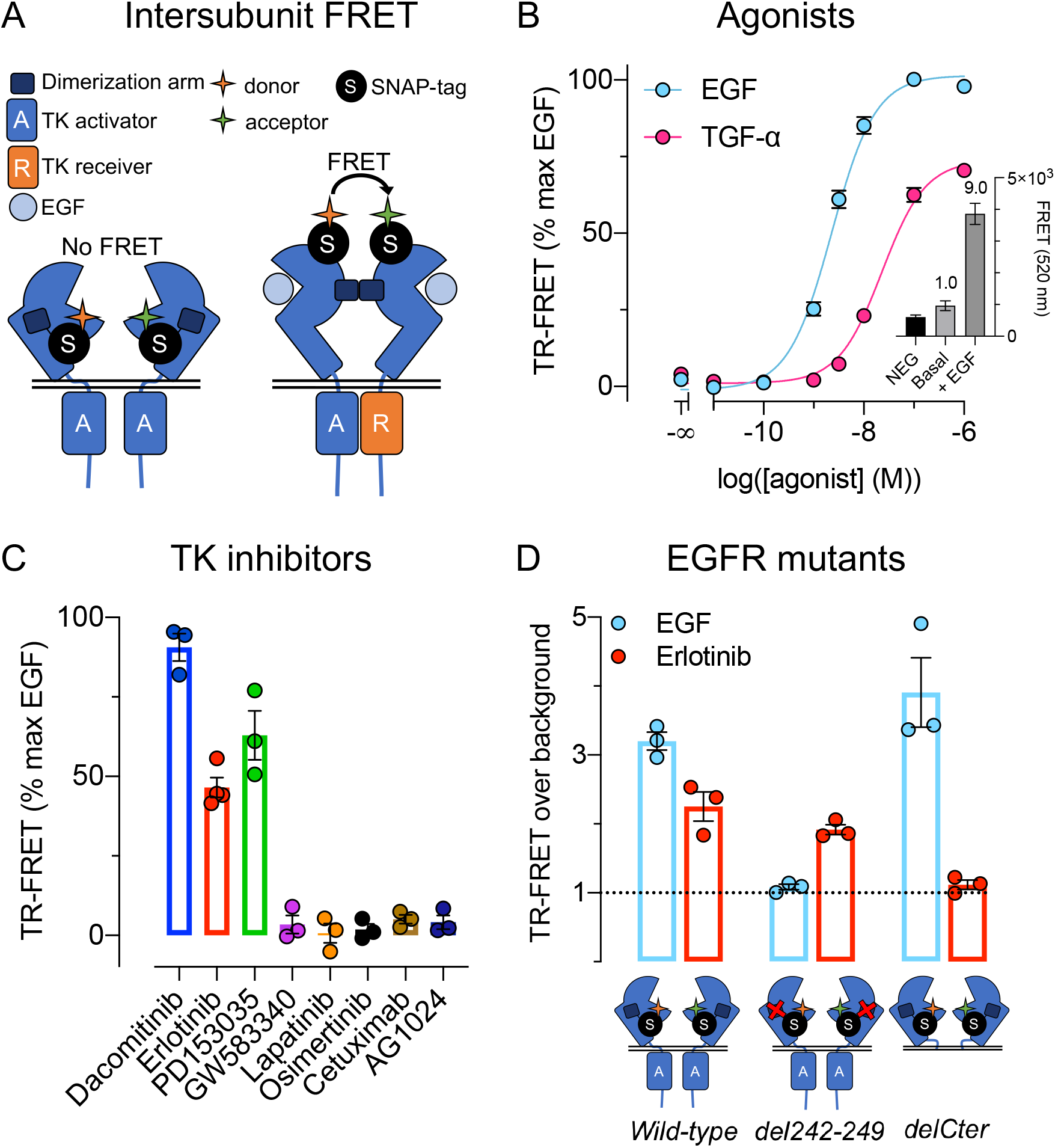
Intersubunit FRET is induced by group I TK inhibitors through the TK domain. *(A)* Cartoon representing a TR-FRET-based EGFR intersubunit FRET assay. EGFR subunits were randomly labelled with 125 nM acceptor and 100 nM donor. *(B)* Intersubunit FRET of EGFR in presence of EGF (blue) or TGF-α (pink). Representative raw FRET values and fold increase of specific FRET are shown in a subgraph. *(C)* EGFR intersubunit FRET induced by saturating concentration of TK inhibitors. *(D)* EGFR intersubunit FRET of *wild-type* EGFR or EGFR mutants induced by 100 nM EGF (blue) or 100 µM erlotinib (red). Data in *B-D* are mean ± SEM of three or more individual experiments.

The FRET between two EGFRs induced by the group I TK inhibitor occurs independently of the dimerization of the extracellular domain, as this effect could still be observed with an EGFR mutant with a disrupted dimerization arm (EGFR_del242-249_) (Fig. 1D and Suppl. Fig. S3C). As expected, the agonists had no effect on this mutant. On the contrary, EGF and not erlotinib induced intersubunit FRET on a receptor deleted of its intracellular domain (EGFR_delCter_) ^17^ (Fig. 1D and Suppl. Fig. S3D). This showed that erlotinib induces intersubunit FRET of the EGFR through the TK domain independently of the dimerization arm.

This proposal was confirmed for the full-length EGFR, when exposure of the dimerization arm was prevented by cetuximab (Fig. 2A). Notably, the TR-FRET signal induced by erlotinib was not affected by cetuximab, while the effect of EGF was fully inhibited (Fig. 2A). As control, we verified that the absence of effect of cetuximab is not due to a specific conformation stabilized by erlotinib that could prevent cetuximab binding. Indeed, cetuximab could still inhibit binding of antibody 58 labelled with d2 (Ab58-d2) in the presence of erlotinib (Fig. 2B, C). These data are consistent with the increase in intersubunit FRET being the result of ligand-induced dimerization rather than of conformational changes of the extracellular domain.

**Fig. 2.**
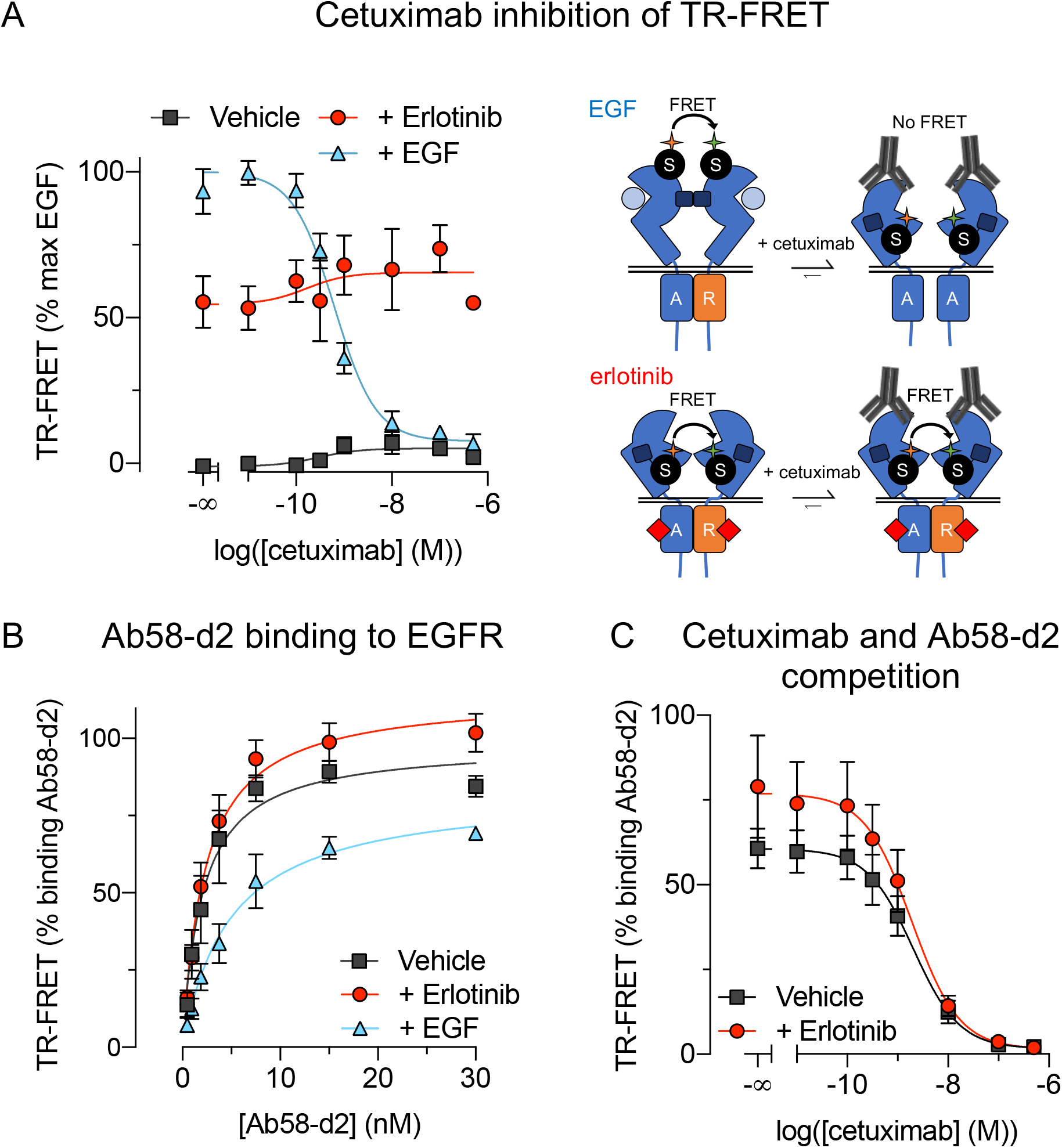
Erlotinib-induced dimers are not inhibited by cetuximab. *(A)* EGFR intersubunit FRET in presence of cetuximab with vehicle (gray), EGF (blue), or erlotinib (red) and a schematic interpretation of the data. *(B)* Binding of Ab58-d2 to SNAP-EGFR that is labelled with 125 nM donor and pre-incubated with saturating concentrations of erlotinib (red) or EGF (blue) or vehicle (gray). *(C)* Binding of Ab58-d2 in presence of cetuximab with (red) or without (gray) erlotinib. Data in *A-C* are mean ± SEM of three or more individual experiments.

### Group I TK inhibitors induce less dimers than agonists

Whether EGF acts by allowing dimer formation, or through conformational changes within pre-formed dimers is still a matter of debate ^15^. The absence of effect of cetuximab, or of the deletion of the dimerization arm on TK inhibitor-induced intersubunit FRET is more consistent with ligand-induced dimerization. To further study this possibility, we examined why TK inhibitors induced a lower maximal FRET signal than agonists. This can be due either to a different conformation of the extracellular part of the EGFR dimer, which would be consistent with pre-formed dimers, or to a lower proportion of receptors in FRET than with agonist. To clarify this point, we analyzed the excited-state lifetime of the sensitized acceptor emission (τ_DA_) as an indication of the conformational state of the receptor, as τ_DA_ is related to the distance between the fluorophores.

As a control for this approach, we analyzed the SNAP-metabotropic glutamate (mGlu) 4 - CLIP-mGlu2 receptor heterodimer, a prototypical preformed dimer both subunits being linked by a disulphide bond ^18^. In that case, a change in FRET cannot be due to a different proportion of dimers, and then only rely on a change in distance due to conformational changes. The C termini of the subunits were modified by addition of the endoplasmic reticulum retention sequences of the GABA_B2_ (C2) and GABA_B1_ (C1) to the respective receptors, so that only heterodimeric receptors were expressed at the cell surface (Suppl. Fig. S4A) ^19^. We labelled each mGlu2-4 heterodimer subunit with SNAP-Lumi4-Tb (donor) and CLIP-Green (acceptor), and measured the FRET signal.

In principle, the excited-state lifetime of the FRET donor emission (τ_D_) is in the millisecond range, and decreases when the donor’s excited state is subject to other deactivation pathways such as FRET. As the τ_DA_ is proportional to the distance between a FRET donor and acceptor, distinct FRET donor-acceptor distances can be determined. The τ_DA_ is measured in a time-resolved manner to discriminate between non-specific and specific signal ^20^.

In the basal mGlu2-4 condition, τ_DA_ was 343 ± 25 µs, corresponding to a high FRET conformation. The τ_DA_ was largely increased to 564 ± 22 µs in the presence of the mGlu2 agonist LY379268, indicating a low FRET conformation (Fig. 3A and B) ^17, 21^. These data confirm that a conformational change within constitutive dimers can be detected by measuring the τ_DA_ values.

**Fig. 3.**
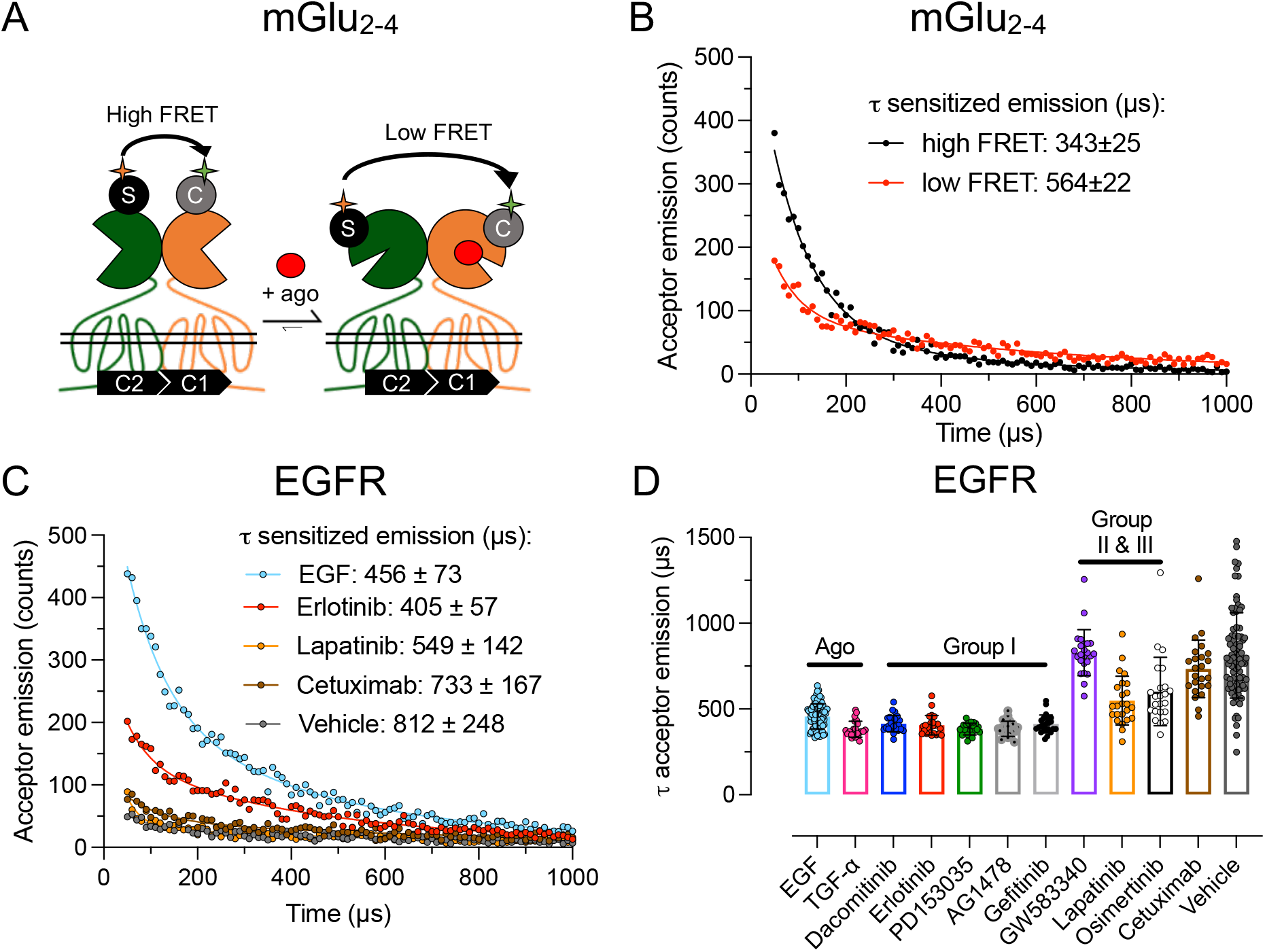
Sensitized emission reveals that higher FRET levels correspond to more receptors in FRET rather than closer proximity of FRET-pair. *(A)* Intersubunit FRET between an mGlu2 (orange) and mGlu4 (green) constitutive dimer labelled with acceptor and donor. *(B)* Representative data of excited-state sensitized acceptor emission of mGlu2-4 intersubunit FRET assay in presence of vehicle (black) and 100 µM LY379268 (red). *(C)* Representative data of excited-state sensitized acceptor emission of EGFR intersubunit FRET assay in presence of 100 nM EGF (blue), 10 µM erlotinib (red), 10 µM lapatinib (orange), 100 nM cetuximab (brown) and vehicle (gray). *(D)* Summary of excited-state sensitized acceptor emission of EGFR intersubunit FRET assay. Data in *D* are individual values ± SD of at least 23 datapoints obtained in at least three individual experiments. Statistical test in *C* is one-way ANOVA with Dunnett’s multiple comparison (Suppl. Table S3).

For the SNAP-EGFR, the τ_DA_ measured in the presence of agonists or group I TK inhibitors were not significantly different (Fig. 3C and D, Suppl. Table S3), despite differences in TR-FRET intensity (Suppl. Fig. 4B). This suggests that the larger TR-FRET level measured with agonists is due to a higher proportion of receptors in FRET, rather than to a distinct donor-acceptor distance. Such data are then consistent with TK inhibitors stabilizing less EGFR dimers than agonists.

### Inhibition of EGF-induced phosphorylation of EGFR and ERK1/2 by TK inhibitors

We then evaluated the efficacy of the inhibitors for EGF-induced phosphorylation of either EGFR or ERK1/2. Phosphorylation of tyrosine residue 1068 of the EGFR (Y1068) and of threonine residue 202 and tyrosine residue 204 of ERK1/2, as detected by the HTRF® signals in an antibody-based sandwich assay (Fig. 4A and Suppl. Fig. 5), are inhibited efficiently by group I, group II TK inhibitors and cetuximab and less efficaciously by group III TK inhibitor osimertinib. Conversely, irrelevant TK inhibitor AG1024 did not inhibit phosphorylation of the EGFR or ERK1/2 (Fig. 4B, C and Suppl. Fig. 5B, D).

**Fig. 4.**
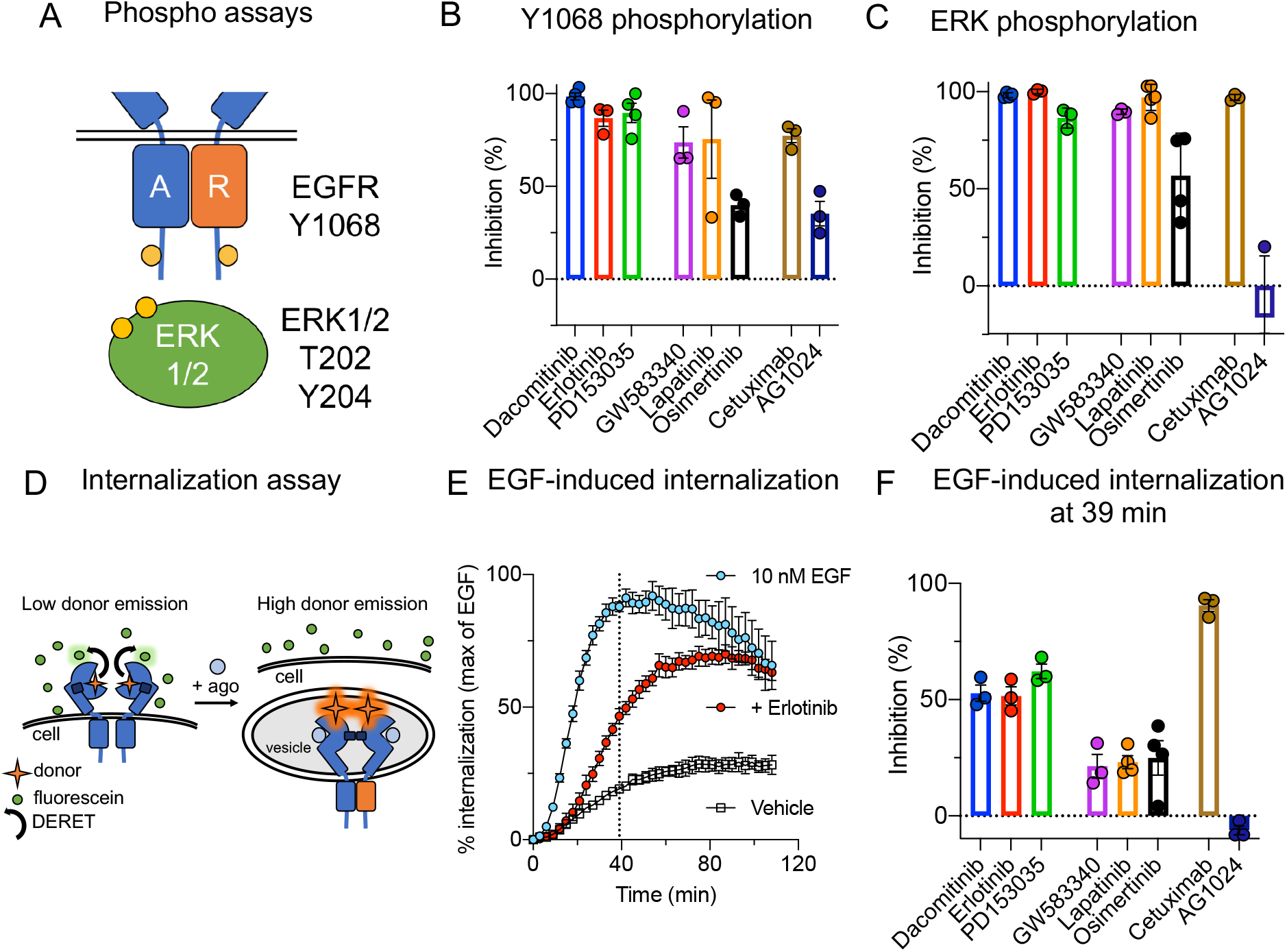
Phosphorylation of EGFR and ERK1/2, and internalization of EGFR. *(A)* Scheme of antibody-based sandwich assays to measure EGFR and ERK1/2 phosphorylation. *(B)* Inhibition of EGF-induced (10 nM) EGFR phosphorylation by 1 µM of dacomitinib, erlotinib, PD153035, lapatinib, osimertinib and AG1024, 0.3 µM GW583340 or 100 nm cetuximab. Inhibitors were pre-incubated for 30 minutes and phosphorylation was measured 30 minutes after stimulation with EGF. *(C)* Inhibition of EGF-induced (10 nM) ERK1/2 phosphorylation by 1 µM of dacomitinib, erlotinib, PD153035, lapatinib, osimertinib and AG1024, 0.3 µM GW583340 or 100 nm cetuximab. Inhibitors were pre-incubated for 30 minutes and phosphorylation was measured 5 minutes after stimulation with EGF. *(D)* Scheme of DERET-based EGFR internalization assay. *(E)* Representative data for EGFR internalization assay in presence of 10 nM EGF (blue), EGF and 1 µM erlotinib (red) or vehicle (gray). *(F)* EGF-induced EGFR internalization in presence of 1 µM of dacomitinib, erlotinib, PD153035, lapatinib, osimertinib and AG1024, 0.3 µM GW583340 or 100 nm cetuximab. Inhibitors were pre-incubated for 2 hours and EGF-internalization was measured after 39 minutes. Data in *B-C* and *F* are mean ± SEM of at least three individual experiments. Data in *E* is represented as mean ± SD.

### TK inhibitors stabilizing dimers slow down EGF-induced EGFR internalization

Then, we setup an internalization assay for the EGFR based on diffusion-enhanced resonance energy transfer (DERET) ^22^. In principle, SNAP-EGFR is labelled with Lumi4-Tb and an excess of fluorescein is added to each well, thereby generating DERET and quenching the Lumi4-Tb emission. Upon internalization of the EGFR, the Lumi4-Tb signal is recovered (Fig. 4D) allowing the detection of EGFR internalization in living cells over time.

We observed a basal internalization in the absence of agonist, but EGF largely increases EGFR internalization, that reaches a plateau after 39 min, followed by a decline, suggesting receptor recycling to the cell surface. Cetuximab fully inhibited EGF-induced internalization but not the basal internalization (Suppl. Fig. 7). Surprisingly, TK inhibitors did not inhibit EGFR internalization (Fig. 4F and Suppl. Fig. 7) as group I TK inhibitors only slowed down EGF-induced EGFR internalization (Fig. 4E), whereas others had no effect (Suppl. Fig. 7).

Despite not inhibiting internalization, it is clear that group I TK inhibitors were more efficacious in slowing down EGFR internalization than the group II and III TK inhibitors and irrelevant TK inhibitor AG1024 (Fig. 4F and Suppl. Fig. 7). We compared internalization of EGFR with fluorescence microscopy, showing comparable results (Suppl. Fig. 6).

The combined data of the 4 assays revealed a different pharmacological profile of each group of compounds for the wild-type EGFR as represented in Figure 5 (Suppl. Fig. S8). Moreover, bias plots for the agonists demonstrated that EGF and TGF-α are more potently inducing internalization and phosphorylation of ERK1/2 (Suppl. Fig. S9). Among the TK inhibitors, only PD153035 more potently inhibited phosphorylation of Y1068 than ERK1/2 (Suppl. Fig. S10).

**Fig. 5.**
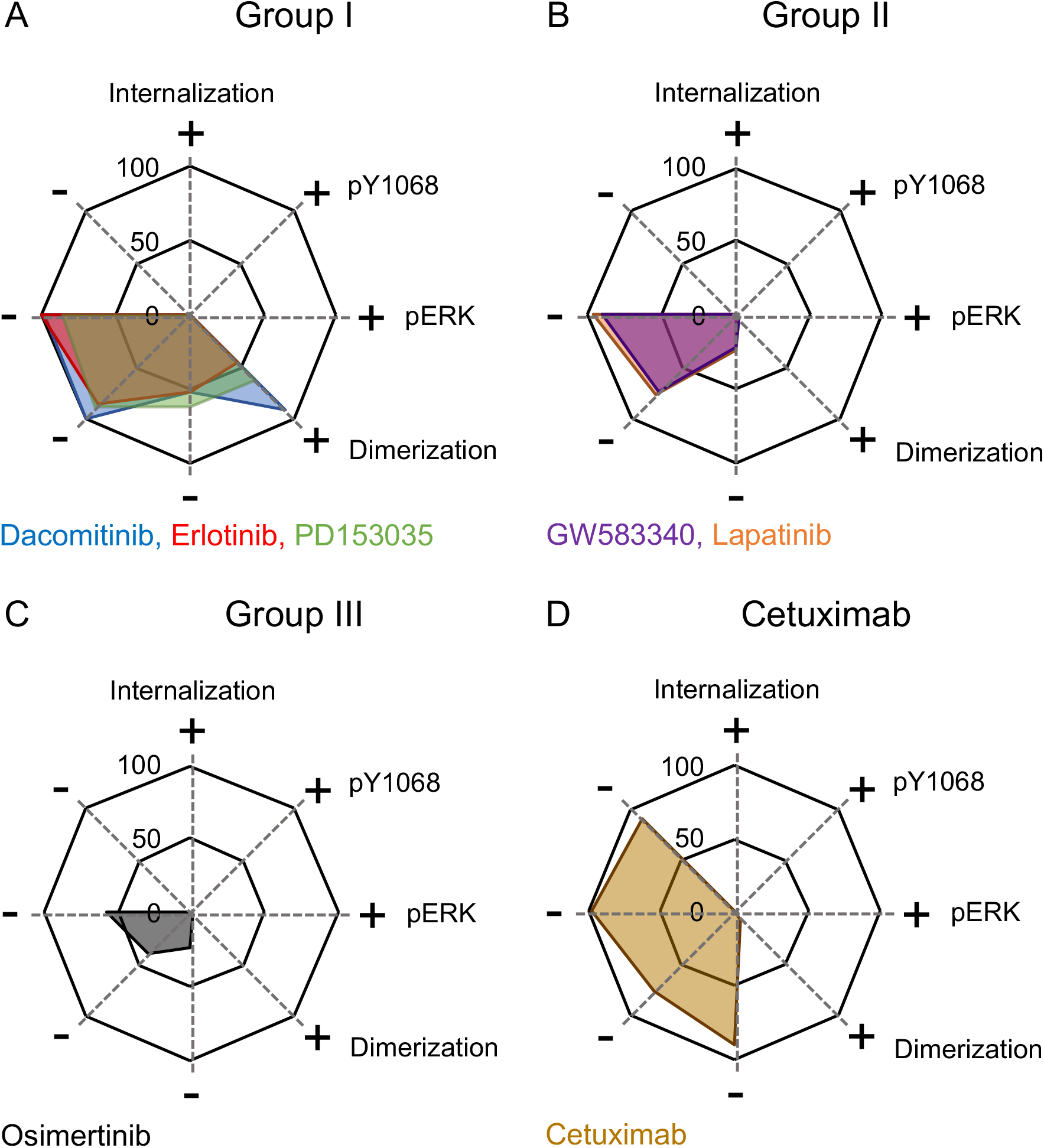
Pharmacological profile of TK inhibitors. *(A)* Group I TK inhibitors dacomitinib (blue), erlotinib (red), PD153035 (green). *(B)* Group II TK inhibitors GW583340 (purple) and lapatinib (orange). *(C)* Group III TK inhibitor osimertinib (gray). *(D)* Cetuximab (brown). All data are from figures 1-4.

### NSCLC-mutated EGFR is insensitive to TK inhibitor-induced dimerization

EGFR_NSCLC_ has an increased activity within the asymmetric dimer ^23^ and it was suggested that the presence of EGFR_NSCLC_ could increase dimer formation ^24^. To investigate if EGFR_NSCLC_ has an increased propensity to induce dimer formation, we developed an intersubunit FRET assay specifically detecting heterodimers of *wild-type* EGFR and EGFR_NSCLC_ or homodimers of EGFR_NSCLC_ (Fig. 6A). For both the heterodimer and the homodimers, EGF induced dimer formation with a similar potency as for *wild-type* EGFR (Fig. 6B).

**Fig. 6.**
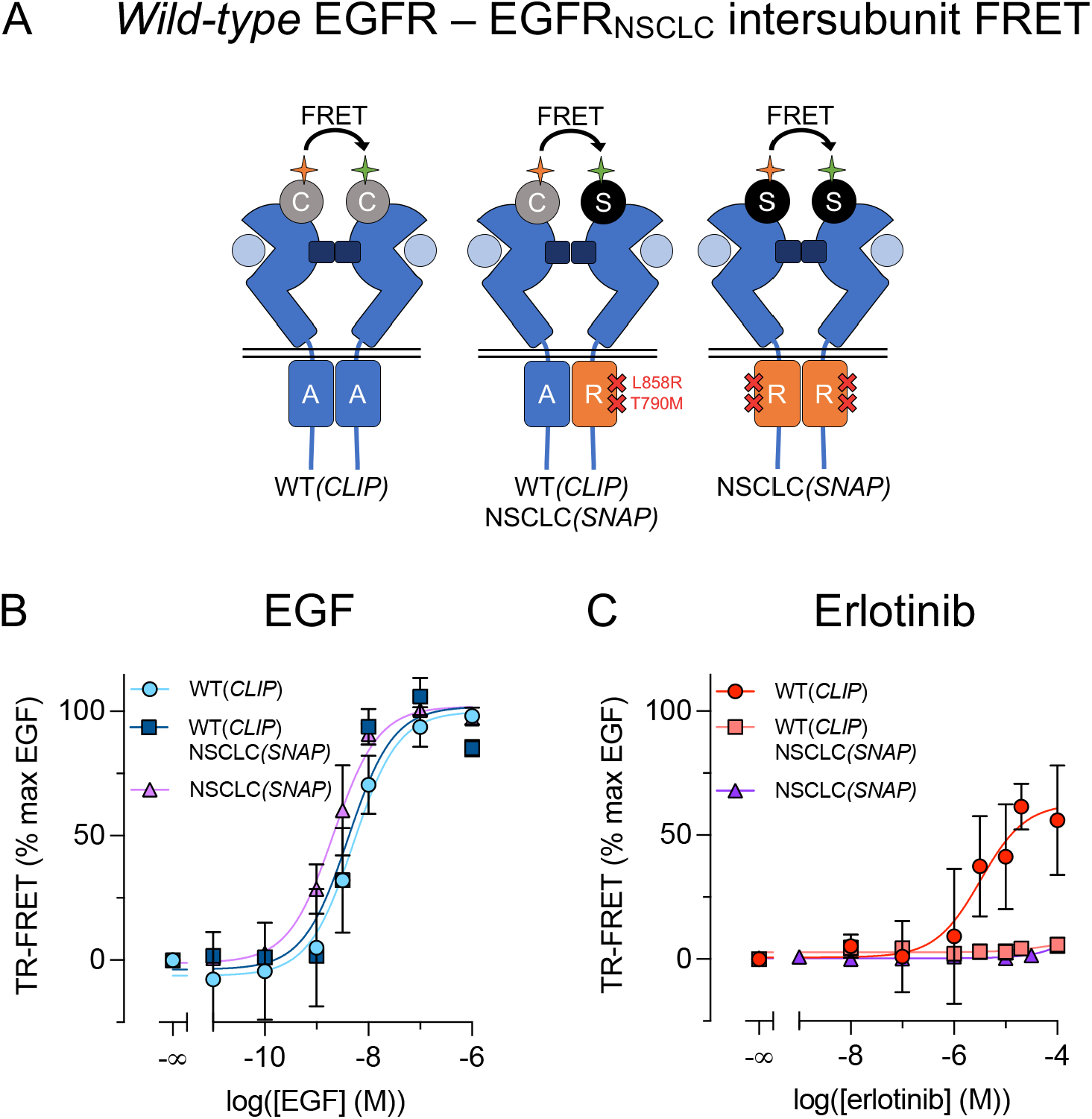
TK domain-induced dimers need binding of two TK inhibitors. *(A)* Scheme of assays to measure intersubunit FRET of the *wild-type* CLIP-EGFR homodimer, CLIP-EGFR + SNAP-EGFR_NSCLC_ heterodimer and SNAP-EGFR homodimer. *(B)* EGF-induced intersubunit FRET. *(C)* Erlotinib-induced intersubunit FRET. Data in *B-C* are represented as mean ± SEM of at least three individual experiments.

As the EGFR_NSCLC_ is resistant to erlotinib-like TK inhibitors due to the T790M substitution in the ATP binding pocket ^5^, we were curious whether erlotinib could induce dimer formation in presence of this mutant. Interestingly, erlotinib could not induce formation of dimers containing EGFR_NSCLC_, either with both subunits mutated, or suggesting that both subunits of an asymmetric TK domain need to be bound by TK inhibitors to induce TK domain dimer formation (Fig. 6C and Suppl. Fig. S11).

## Discussion

Although of clinical importance, several aspects of EGFR activation and the mode of action of various TK inhibitors remain unclear. This study brings clear evidence for the agonist-induced dimer formation model of EGFR activation. It also reveals that formation of dimers can be induced by some TK inhibitors through interaction of the TK domains, resulting from a specific conformation of the TK dimer. Eventually, our data show that EGF-induced internalization is mainly driven by a specific conformation of the EGFR dimer, and not by receptor activity, allowing some, but not all TK inhibitors to slow down this process.

Despite years of research on EGFR two models of receptor activation were still discussed ^15^. One model proposes that agonists like EGF act by promoting dimer formation ^25^, while the other proposes that agonists stabilize a specific conformation of a pre-formed EGFR dimer ^26^. Our data strongly support the first model. Indeed, through FRET measurements between N-terminal tags, with cell surface receptors labelled exclusively, we detected a very low FRET signal under basal condition in a cell line stably expressing SNAP-EGFR (Suppl. Fig. S1). This signal is largely increased (9-fold) in the presence of agonist (Fig. 1B). The low FRET under basal condition is unlikely due to pre-formed dimers with a conformation leading to a large distance between the N-termini carrying the SNAP-tags, for two main reasons. First, although a very low FRET was measured under the basal condition, the estimated ¹ _DA_ is only twice of that measured with the active dimer (Fig. 3C), consistent with the low FRET resulting mainly from a very low number of dimers. Such finding is also consistent with the distances between the N-termini of the proposed inactive and active dimers ^24^. Second, when dimers were stabilized through their TK domains using some TK inhibitors, the extracellular domains do not appear to interact via the dimerization arm, and are then likely in an inactive conformation possibly similar to that proposed by others for the pre-formed dimers ^24^. If so, such pre-formed dimers should generate a FRET signal with a ¹ _DA_ similar to that obtained with the EGF bound EGFR dimers. Accordingly, our data are consistent with a very small proportion of the EGFR subunits at the cell surface involved in FRET, either because of a low proportion of pre-formed dimers, or because of random collisions of EGFR monomers (by-stander FRET).

Consistent with previous studies ^27-29^, some TK inhibitors can also induce dimer formation. Because the dimerization arm is not necessary, it suggests a main role of the TK domain interaction in this process. It is interesting to note that TK inhibitors inducing dimer formation can bind either to the Activator-like or the Receiver like TK domain ^30^ (Fig 7), thereby allowing the formation of asymmetric TK domain dimers, as expected in the active form of the EGFR dimer ^31^. The proportion of the Activator and Receiver forms may then dictate the number of possible dimers – i.e. the higher the proportion of one species (i.e. Activator or Receiver), the lower the number of possible Activator-Receiver dimers. This explains why TK inhibitors are not being able to promote the formation of the same amounts of dimers as EGF (Suppl. Fig. S4). In contrast, TK inhibitors stabilizing a specific conformation of the TK domain would not favor dimer formation according to this model, in agreement with what observed here. This reveals the critical importance of the effect of TK inhibitors on the TK domain conformation on their capacity to stabilize EGFR dimers.

**Fig. 7.**
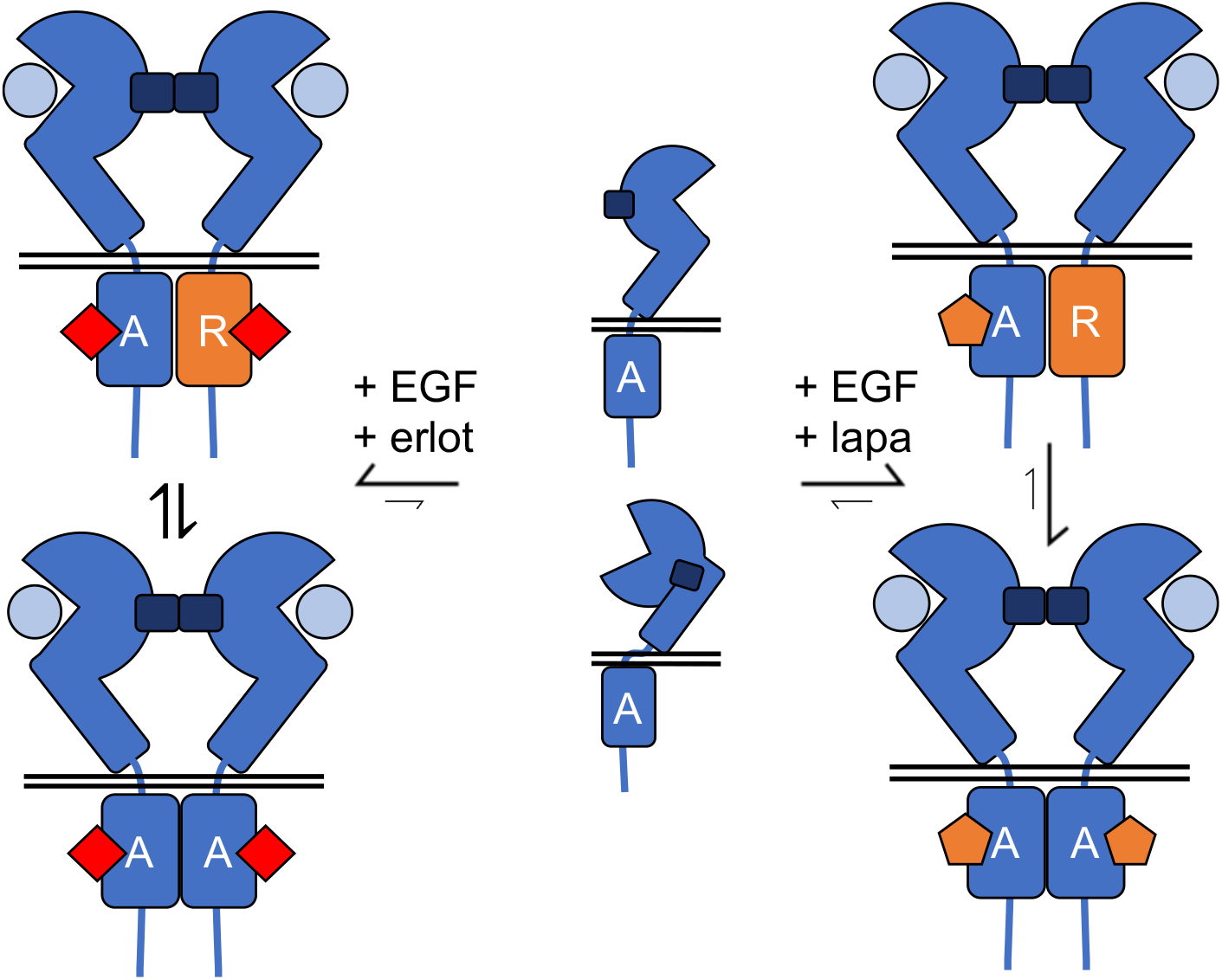
Schematic hypothesis of stabilized dimers by group I and group II TK inhibitors in presence of EGF. Group I TK inhibitors stabilize both Activator and Receiver conformations and group II TK inhibitors only Activator conformations of the TK domain. In presence of erlotinib there are more inactive Activator-Receiver dimers than in presence of lapatinib.

As already reported ^32^, EGFR can rapidly engage into internalization upon agonist activation, though the number of internalized receptors rapidly declined after a peak, possibly due to receptor recycling to the cell surface. Such a process has been known to limit EGFR signaling and can involves clathrin coated pits^33-36^. Internalized EGFR have also been shown to still generate specific signals before most of them are being degraded^33, 34^. This agonist-induced EGFR internalization has soon been assumed to result from EGFR activity ^34, 37^. Indeed, deletion of the TK domain, or the M721 mutation that fully inhibits TK activity prevent agonist-induced EGFR internalization ^34, 37^. Surprisingly, we show here that this process is not prevented by any of the TK inhibitors tested despite their full inhibitory effect on agonist-induced receptor phosphorylation and ERK activation. As such, the internalization process may likely be the result of a specific conformation of the dimer. Such a hypothesis is still compatible with the absence of internalization of a TK domain deleted receptor. As for the M721 mutation that also prevents internalization, it remains possible that this mutation also affects the general conformation of the TK dimer within the activated receptor, but this remains to be clarified. Interestingly, the differential effect of TK inhibitors -those not promoting dimers having no effect on the internalization process while those promoting dimers slow down this process -highlights an essential role of the TK dimer conformation. However, it is surprising to see that TK inhibitors promoting EGFR dimer formation, do not promote internalization.

To explain these apparently contradictory results, one should consider the possible conformation of the intracellular part of the receptor. Indeed, TK inhibitors that stabilize the Activator state of the TK domain, such as lapatinib, do not promote dimer formation, and do not affect EGF-induced internalization (Fig 7). This suggests that a symmetric EGFR dimer in which both TK domains are in the inactive Activator state, are perfectly prone to internalization after direct association of the extracellular domains. In contrast, TK inhibitors like erlotinib that can stabilize either the Activator or the Receiver state (Fig 7), promote dimer formation likely through a stable asymmetric TK domain dimer composed of an Activator and Receiver TK domain. They do not induce receptor internalization, while they only slow down EGF-induced internalization. As such, one is tempted to propose that the asymmetric Activator-Receiver dimer is not prone to internalization. Such hypothesis explains our results, but would certainly need further support to be fully validated.

Previously, agonist-induced conformational models of the EGFR ^38, 39^ and the role of agonist binding kinetics in ligand bias have been described ^40, 41^. For TK inhibitors, such studies are less well known, whereas it could improve understanding of their mode of action. Generation of pharmacological profiles and bias plots has recently been proposed as mode to improve EGFR drug design ^42^. Techniques for determining pharmacological profiles or ligand bias are more common in the research field of G protein-coupled receptors (GPCRs) and contributing to GPCRs being the most targeted family of receptors by drugs on the market ^43, 44^. In this study, such techniques have been adapted for an analysis of the EGFR signaling.

Efforts to expose functional selectivity induced by EGFR ligands indicated that agonists have different signaling kinetics by stabilizing different conformations of the extracellular domain ^38^. Bias plots for agonists EGF and TGF-α reveal that they are more potent for phosphorylation of ERK and internalization than dimerization and phosphorylation of Y1068 (Suppl. Fig. S9). Moreover, we found that most TK inhibitors inhibit phosphorylation of Y1068 and ERK with similar potency, except for PD153035, which more potently inhibits phosphorylation of Y1068 (Suppl. Fig. S10C).

Some TK inhibitors may impact EGF binding through allosteric modulation of the EGFR conformation ^39, 45^. We show that TK inhibitors stabilize distinct dimers, resulting in altered EGFR trafficking. Previous reports linked dimerization to improved cellular survival ^28^ and decreased efficacy for some TK inhibitors ^46^. Moreover, disruption of dimerization could be an antitumoral mechanism as well ^47^, confirming its significance. The role of internalization in tumor survival is not fully clear, nonetheless it has been suggested that *wild-type* NSCLC patients (i.e. no resistance mutations) could benefit from blocking clathrin-mediated endocytosis of EGFR ^48, 49^. This implies that ligands that reduce EGFR internalization (i.e. dacomitinib, erlotinib, PD153035) could induce positive outcomes for *wild-type* EGFR in NSCLC patients.

To our best knowledge there are no reports of TK inhibitors inducing dimerization of the EGFR_NSCLC_ or heterodimers of *wild-type* EGFR and EGFR_NSCLC_. In our model we do not observe pre-formed homo-and heterodimers containing EGFR_NSCLC_, as the dimers are mainly observed upon EGF or TGF-α activation. The potencies for dimerization are not significantly different from *wild-type* EGFR (Suppl. Table S2). None of the tested TK inhibitors induces dimerization, suggesting there is a direct or indirect loss of potency for the heterodimer due to altered binding (45, 46) or increased affinity for ATP ^50^. In tumor cells of NSCLC patients, different populations of EGFR heterodimers could exist, among them *wild-type* EGFR-EGFRNSCLC ^5^and EGFR-ErbB2 heterodimers ^51^. Investigations on the effect of drugs on these heterodimeric receptors, could help improving treatment strategy as they may function as additional drug targets. Another approach could be the use of allosteric modulators to decrease off-target effects ^7^. The use of allosteric compounds like EAI045 for treating NSCLC has recently been reviewed ^2^.

Overall, this study reveals the importance of the conformational state of the TK domains within the EGFR dimer in signaling and trafficking. As TK inhibitors have various effect on such conformations, this may explain their biased effects on dimerization and internalization, properties that likely have some importance in the cellular physiology of EGFRs.

## Materials and methods

Reagents and the protocols for cell culture, cell preparation for experiments, generation of SNAP-EGFR_NSCLC_ and CLIP-EGFR plasmids, lipofectamine transfection for experiments, generation of HEK293 cells stably expressing SNAP-EGFR and SNAP-EGFR_NSCLC_, intersubunit FRET, binding and displacement of Ab58-d2, ERK1/2 phosphorylation, phosphorylation of tyrosine residue 1068 of the EGFR (Y1068), internalization, fluorescence microscopy and data analysis can be found in the supplementary information. Monoclonal stable cell lines stably expressing SNAP-EGFR or SNAP-EGFR_NSCLC_ were generated by lipofectamine transfection. Positive cells were selected by G418 and single cells were sorted with fluorescence-activated cell sorting. An estimated 256,000 individual SNAP-EGFRs (i.e. 4-fold lower than A341 squamous carcinoma) are present on the cell surface and expression levels were stable over time (Suppl. Fig. S1).

### Excited state-lifetime of sensitized acceptor emission to determine receptor conformations

At 24 hours after lipofectamine transfection (for SNAP-mGlu4-C2-KKXX and CLIP-mGlu2-C1-KKXX constructs) or transfer of cells stably expressing SNAP-EGFR into a 96-wells plate, medium was replaced by ice-cold DMEM containing SNAP-Lumi4-Tb (100 nM) and SNAP-Green (125 nM) for SNAP-labelling and CLIP-Lumi4-Tb (1 µM) and CLIP-Green (1 µM) for CLIP-labelling. Cells were incubated for 90 minutes at 4 °C and carefully washed four times with ice-cold Tag-Lite buffer. LY379268 (100 µM), EGF (100 nM), TGF-α (100 nM), dacomitinib (10 µM), erlotinib (10 µM), PD153035 (10 µM), AG1478 (10 µM), GW583340 (5 µM), lapatinib (10 µM), osimertinib (10 µM), cetuximab (10 nM) or vehicle was incubated in ice-cold Tag-Lite buffer for 30 minutes at 4 °C. Luminescence decay at 520 nm was measured after 150 flashes/well with the UV-pulsed nitrogen laser (337 nm) of the PHERAstar FS microplate reader. Decay was measured from 50 to 5000 µs and was fitted using the biexponential decay function in GraphPad Prism software (version 9.0.1.), which is the preferred model in an extra sum-of-squares F-test compared to a mono-exponential decay function. The excited-state lifetime of the sensitized acceptor emission (τ_DA_) was calculated with a least-squares fit. The apparent amplitude of slow (A_s_) and fast (A_f_) components of the biexponential decay may vary due to adaptation to multiple conformations or interactions with the antenna, increasing the complexity of its decay ^52^. Typically, the apparent A_f_ was larger than A_s_ whereas this value is likely overestimated and should be corrected ^20^. The true fraction of the slow decay species (α_DA_) is based on the resonance energy transfer rate constant, as described by Heyduk *et al*.^*20*^. After applying this correction, α_DA_ is in the range of 0.75-0.88 for all conditions (Suppl. Table S1).

## ACKNOWLEDGEMENTS

We thank the ARPEGE platform facilities at the Institut de Génomique Fonctionnelle for all fluorescence-based assays and the MRI platform facilities at the Institut de Génétique Humaine for FACS and microscopy assays. We thank dr. Robert Quast for discussions. J.-P.P. was supported by la Fondation pour la Recherche Médicale (ref. DEQ20170336747), Cisbio Bioassays (EIDOS collaborative team IGF-CISBIO, ref. 039293), the Fond Unique Interministériel of the French government (FUI, Cell2Lead project), the Agence Nationale pour la Recherche (ANR 18-CE11-0004-01) and LabEx MAbImprove (ref. NR-10-LABX-5301). J.-P.P. and L.P. were further supported by the Centre National de la Recherche Scientifique and the Institut National de la Santé et de la Recherche Médicale. J.H. was supported by a fellowship from la Région Occitanie and Cisbio Bioassays (TransACT, ref. 156544) and la Ligue contre le cancer (PhD grant, ref. IP/SC-16487). *Add the financial support from J*.*M*.*Z. and F*.*C*.*-S*.

## Notes

### Competing Interest Statement

JP Pin's laboratory is connected to Perkin-Elmer/Cisbio, through the common laboratory Eidos.

