## Supplementary information for "Different conformations of EGF-induced receptor dimers involved in signaling and internalization"

**Reagents**

CLIP-Lumi4Tb (cat. no. SCLPTBF), CLIP-Green (cat. no. SCLPGRNF), SNAP-Lumi4Tb (cat. no. SSNPTBX), SNAP-Green (cat. no. SSNPGRNZ), SNAP-Red (cat. no. SSNPREDZ), Tag-Lite® buffer (cat. no. LABMED), HEK293 cells (cat. no. 300192, CLS Cell Lines Service GmbH, Germany) and the plasmids for SNAP-EGFR_delCter_ and SNAP-EGFR_del242-249_ were provided by Cisbio Bioassays. LY379268 (cat. no. 2453) was bought from Tocris Bioscience. AG1478 (cat. no. T4182), EGF (cat. no. SRP3196), TGF- α (cat. no. T7924), dacomitinib (cat. no. PZ0330), PD153035 (cat. no. SML0564), gefitinib (cat. no. SML1657) GW583340 (cat. no. G3545) and lapatinib (cat. no. SML2259) were bought from Sigma-Aldrich. Erlotinib (cat. no. S7786) and cetuximab (cat. no. A2000) were bought from Selleckchem. Osimertinib (cat. no. I-9701) was bought from Achemblock. AG1024 (cat. no. 121767) was bought from Calbiochem.

**Cell culture**

HEK293 (CLS) cells and HEK293 cells stably expressing SNAP-EGFR and SNAP-EGFR_L858R+T790M_ were cultured in DMEM medium supplemented with 10% fetal calf serum (cat. no. F2442, Sigma-Aldrich), 50 IU/mL penicillin (P4333, Sigma-Aldrich) and 50 µg/mL streptomycin (P4333, Sigma-Aldrich), 1% non-essential amino acids (11140-035, Invitrogen) and 2 mM HEPES (EGFR medium) at 37° C and 5% CO_2_. Cells were grown in P150 petri dishes and cell splitting was performed every two or three days. The culture medium was removed by aspiration and cells were washed with 10 mL PBS. Cells were detached by incubating them in 3 mL of Trypsin-EDTA (0,05%) (cat. no. 25300054, ThermoFisher Scientific) for 3 minutes at 37 °C and 5% CO_2_. Cells were then diluted to the desired concentration and added to new P150 petri dishes with fresh culture medium.

**Cell preparation for experiments**

A black 96-well plate was coated with of poly-L-ornithine (ref. P4957, Sigma-Aldrich) by incubating for 30 minutes at 37 °C. After 30 minutes, wells were washed with PBS to remove any non-bound poly-L-ornithine (PLO). HEK293 cells were grown in EGFR medium at 37 °C and 5% CO_2_ in P150 petri dishes till 80% confluency. To detach cells, they were incubated for 3 minutes at 37 °C and 5% CO_2_ in 3 mL 0.05% trypsin-EDTA. Trypsin was inactivated by addition of an excess of EGFR medium and cells were centrifuged and added to a 96-well plate at a concentration of 5,000, 50,000 or 100,000 cells/well in EGFR medium depending on the experiment and incubated for 24 hours at 37 °C and 5% CO_2_.

**Generation of non-small cell lung cancer mutated-EGFR and CLIP-EGFR plasmids**

The SNAP-EGFR_NSCLC_ and CLIP-EGFR plasmids were generated by modifying the previously described SNAP-EGFR plasmid (1). To generate the two single-point mutations (i.e. L858R and T790M) in the SNAP-EGFR construct, firstly the L858R point mutation was generated in SNAP-EGFR to obtain SNAP-EGFR_L858R_-plasmid. Next, a T790M point mutation was generated in the EGFR_L858R_-plasmid to obtain the SNAP-EGFR_L858R+T790M_-plasmid (SNAP-EGFR_NSCLC_). Corresponding sense and antisense primers were bought from Eurofins Genomics and introduced in the SNAP-EGFR by site-directed mutagenesis following the manufacturer’s protocol (Agilent). For the generation of the CLIP-EGFR plasmid a gene containing the CLIP-tag was generated and substituted for the SNAP-tag by GeneCust through the generation of a synthetic gene.

**Lipofectamine transfection for experiments (for transient receptor transfection)**

Cells were prepared following the ‘’cell preparation for experiments’’ protocol. Transient plasmid transfection was done following the reverse lipofectamine transfection manufacturer’s (Lipofectamine™ 2000, Invitrogen). In short, a DNA mixture of 150 ng pcDNA (i. 4 ng of SNAP-EGFR, ii. 100 ng of CLIP-EGFR or iii. 40 ng SNAP-mGlu4-C2 and 80 ng CLIP-mGlu2-C1 qsp 150 ng with empty pRK6 vector) and 0.375 µL Lipofectamine 2000™ reagent qsp 50 µL OptiMEM (cat. no. 31985062, Thermo Fisher) per well was prepared. The prepared DNA mixture was added to HEK293 cells or a monoclonal HEK293 stable cell line in a 96-well plate depending on the experiment. The cell/DNA mixture was then incubated for 24 hours at 37 °C and 5% CO_2_.

**Generation of HEK293 cells stably expressing SNAP-EGFR and SNAP-EGFR_NSCLC_**

We incubated 250,000 HEK293 cells per well in DMEM supplemented with 10% FBS in a 6 wells plate for cell culture for 24 hours at 37 °C and 5% CO_2_. After 24 hours we removed the medium by aspiration and added 1 mL of DMEM supplemented with 10% FBS and 1 mL OptiMEM containing 3,75 µL lipofectamine 2000™ reagents and 1500 ng DNA (i.e. 40 ng of the plasmid coding for SNAP-EGFR and 1460 ng of the plasmid coding for an empty DNA vector). Cell splitting was performed when confluency reached 80% and medium was refreshed every 2 or 3 days. At day 4 after lipofectamine transfection, medium was replaced by DMEM supplemented with 10% FBS and 0.3 mg/mL G418 (selection medium), a neomycin analog, to start selection as the used vector contains neomycin resistance. Cells were grown in 6-wells plates and at day 16 after transfection, cells were transferred into P100 plates and further grown in selection medium. At day 26 after transfection, cells were labelled at 16 °C for 1 hour with SNAP-Green and washed to remove unbound SNAP-Green. Next, cells were released from the plates with enzyme-free dissociation buffer (cat. No. 13151014, Gibco™) and diluted in PBS to a concentration of less than 1 million cells/mL. Single cell sorting was performed by the BD FACSMelody™ Cell Sorter based on signal at 488 nm, corresponding to the emission spectrum of SNAP-Green. Before cell sorting, we set the negative gate according to the counts measured for non-transfected HEK293 cells incubated with SNAP-Green. Cells that were positive were sorted in clear-bottomed black 96-wells plates and grown in selection medium. Positive clones were identified and transferred to clear 24-wells plates 40 days after transfection and 6-wells plates 45 days after transfection. After 46 days of transfection, cells were transferred to P100 plates. At 48 days after transfection, expression levels were regularly verified.

**Intersubunit FRET assay**

For the measurement of the intersubunit FRET, 50,000 monoclonal HEK293 cells stably expressing SNAP-EGFR or SNAP-EGFR_NSCLC_ were incubated at 37 °C and 5% CO_2_ in a black PLO-coated 96-wells plate 24 hours prior to the start of the experiment. EGFR medium was replaced 90 minutes before the start of the experiment by 50 µL cold serum-free DMEM supplemented with 100 nM SNAP-Lumi4-Tb and 125 nM SNAP-Green for the SNAP-SNAP-labelling or 125 nM SNAP-Green and 1 µM CLIP-Lumi4-Tb for SNAP-CLIP-labelling and cells were stored at 4 °C. Cells were then washed four times with Tag-lite buffer and 50 µL Tag-Lite containing ligands were added. Cetuximab was always pre-incubated 30 minutes before addition of the other compounds. After addition of the ligands, the cells were incubated at 37 °C and 5% CO_2_ for 30 minutes. Next, the plates were read in the PHERAstar FS microplate reader with 30 flashes/well of a 337 nm UV-pulsed nitrogen laser. Emission levels at 520 nm and 620 nm were integrated between 50 and 450 µs. The HTRF®-ratio was then calculated as follows:

$HTRF®-ratio=\frac{\int_{50}^{450} 520 nm signal}{\int_{50}^{450} 620 nm signal}*10,000$.

Curve fitting was performed with GraphPad Prism software using nonlinear regression log(agonist) vs. response (three parameters) analysis. Values were normalized to the maximum and minimum response of EGF.

**Binding and displacement of Ab58-d2**

For the measurement of the binding of Ab58-d2 to SNAP-EGFR, 50,000 monoclonal HEK293 cells stably expressing SNAP-EGFR were incubated at 37 °C and 5% CO_2_ in a black PLO-coated 96-wells plate 24 hours prior to the start of the experiment. EGFR medium was replaced by cold serum-free DMEM supplemented with 100 nM SNAP-Lumi4-Tb 90 minutes prior to the experiment. Cells were then washed four times with Tag-lite buffer. Then, vehicle, 10 µM erlotinib or 10 nM EGF and Ab58-d2 were added to the cells for binding of Ab58-d2. For the displacement of Ab58-d2, vehicle or 10 µM erlotinib, 3.75 nM Ab58-d2 and cetuximab were added to the cells. The cells were incubated for 30 minutes at 37 °C and 5% CO_2._ After incubation, the 96-well plate was read in the PHERAstar FS microplate reader using a 30 flashes/well with a 337 nm UV-pulsed nitrogen laser. Emission levels at 665 nm and 620 nm were integrated between 50 and 450 µs. The HTRF®-ratio was then calculated as follows:

$HTRF®-ratio=\frac{\int_{50}^{450} 665 nm signal}{\int_{50}^{450} 620 nm signal}*10,000$.

**ERK1/2 phosphorylation**

For measuring phosphorylation and total ERK1/2 we added 5,000 monoclonal HEK293 cells stably expressing SNAP-EGFR per well to a black PLO-coated 96-well plate 24 hours prior to starting the experiment. EGFR medium was replaced by 50 µL pre-heated serum-free DMEM 3 hours prior to the experiment and cells were kept at 37 °C and 5% CO_2_. Ligands were prepared in serum-free DMEM and heated to 37 °C. Inhibitors were pre-incubated for 10 minutes after which EGF was added. After 5 minutes of stimulation with EGF, the medium was removed by plate flicking and cells were lysed in 50 µL lysis buffer directly. After cells were lysed, 16 µL of lysate was transferred to a white 384-well plate and 4 µL of an anti-phospho-ERK1/2 antibody mixture (cat. no. 64AERPEH, Cisbio Bioassays) was added to the lysate. During the optimization, we determined that total ERK1/2 levels were not influenced by any of the ligands and therefore 16 µL of lysate of cells stimulated with vehicle was transferred to a 384-wells plate and 4 µL of an anti-ERK1/2 antibody mixture (cat. no. 64NRKPEH, Cisbio Bioassays) was added to the lysate. Both antibody mixtures consist of antibodies labelled with Eu^3+^-cryptate and d2, TR-FRET-compatible fluorophores. The lysate and antibody mixtures were incubated for minimally 4 hours at room temperature. After incubation, 384-wells plates were read in the PHERAstar FS microplate reader using 30 flashes/well with a 337 nm UV-pulsed nitrogen laser. Emission levels at 665 nm and 620 nm were integrated between 50 and 450 µs. The HTRF®-ratio was then calculated as follows:

$HTRF®-ratio=\frac{\int_{50}^{450} 665 nm signal}{\int_{50}^{450} 620 nm signal}*10,000$.

For each sample, the HTRF®-ratios of the phospho-ERK1/2 and total ERK1/2 were divided to calculate the signal over noise ratio (S/N):

$S/N= \frac{HTRF®-ratio (phospho ERK1/2)}{HTRF®-ratio (total ERK1/2)}$.

S/N values were plotted as a negative logarithmic function of ligand concentrations and a curve was fitted with GraphPad Prism software, using a log(agonist) vs. response (three parameters) analysis for agonists and log(inhibitor) vs. response – response (three parameters) analysis for inhibitors. Values were normalized and the bottom of the curve of EGF was set as 0% and the top of the curve of EGF was set at 100%. The best fit values of the logEC50 and logIC50 values were determined by performing a least-squares fit.

**Phosphorylation of Y1068 of the EGFR**

The phosphorylation and total EGFR were measured by adding 50,000 polyclonal HEK293 cells expressing SNAP-EGFR per well to a black PLO-coated 96-well plate 24 hours prior to starting the experiment. EGFR medium was replaced by 50 µL pre-heated serum-free DMEM 3 hours prior to the experiment and cells were kept at 37 °C and 5% CO_2_. Ligands were prepared in serum-free DMEM and heated to 37 °C. Inhibitors were pre-incubated for 30 minutes after which EGF was added. After 30 minutes of stimulation with EGF, the medium was removed by plate flicking and cells were lysed in 50 µL lysis buffer directly. After cells were lysed, 16 µL of lysate was transferred to a white 384-well plate and 4 µL of an anti-phospho-Y1068-EGFR antibody mixture was added to the lysate. During the optimization, we determined that total EGFR levels were not influenced by any of the ligands. The antibody mixture consists of antibodies labelled with Eu^3+^-cryptate and d2. The lysate and antibody mixture were incubated for minimally 4 hours at room temperature. After incubation, 384-wells plates were read in the PHERAstar FS microplate reader using 30 flashes/well of a 337 nm UV-pulsed nitrogen laser. Emission levels at 665 nm and 620 nm were integrated between 50 and 450 µs. The HTRF®-ratio was then calculated as follows:

$$HTRF®-ratio=\frac{\int_{50}^{450} 665 nm signal}{\int_{50}^{450} 620 nm signal}*10,000$$

HTRF®-ratio values were plotted as a negative logarithmic function of ligand concentrations and a curve was fitted with GraphPad Prism 9.0 software (ref.), using a log(agonist) vs. response (three parameters) analysis for agonists and log(inhibitor) vs. response (three parameters) analysis for inhibitors. Values were normalized and the bottom of the curve of EGF was set as 0% and the top of the curve of EGF was set at 100%. The best fit values of the logEC50 and logIC50 values were determined by performing a least-squares fit.

**Internalization**

To measure internalization of the EGFR, we adapted an internalization assay previously described for GPCRs labelled with a SNAP-tag (2). We added 100,000 monoclonal HEK293 cells stably expressing SNAP-EGFR in EGFR medium per well to a flat-bottomed white PLO-coated 96-well plate 24 hours prior to starting the experiment and were kept at 37 °C and 5% CO_2_. After removal of the EGFR medium by aspiration, cold 100 nM SNAP-Lumi4Tb in serum-free DMEM was added and incubated at 4 °C during 90 minutes. Labelling of the receptors at the cell surface was performed at 4 °C to prevent constitutive internalization. Ligands were prepared in a 25 µM fluorescein solution in Tag-lite buffer (FB) and kept on ice. After 90 minutes, plates were washed three times with Tag-lite buffer and inhibitors in pre-cooled FB were added to the plate and incubated for 30 or 120 minutes. After incubation, agonists (i.e. EGF or TGF-α) in pre-cooled FB were added to the plate and the plate was put in the PHERAstar FS microplate reader that was pre-heated at 37 °C and measurements were started. Each 3 minutes for 120 minutes, cells were excited by a UV-pulsed nitrogen laser at 337 nm and emission spectra at 620 nm between 1500 and 3000 µs and 520 nm emission spectra between 160 and 560 µs were integrated. HTRF®-ratio was calculated as follows:

$$HTRF®-ratio= \frac{\int_{1500}^{3000} 620 signal}{\int_{160}^{560} 520 signal}*10,000$$

HTRF®-ratio values for agonists were plotted as a negative logarithmic function of ligand concentrations and a curve was fitted with GraphPad Prism 9.0 software (ref.), using a log(agonist) vs. response three parameters analysis. The bottom of the curve of EGF was set as 0% and the top of the curve of EGF was set at 100%. The best fits of the logEC50 and logIC50 values were determined by a least-squares fit.

To determine the slowing down of the rate of internalization, values were normalized and the HTRF®-ratio at time-point 0 was set as 0% and the top of the peak value of the response to 100 nM of EGF (typically after 30 to 39 minutes) was set as 100%. We then used the normalized value of inhibitor in presence of 10 nM EGF at the time point of the peak of 100 nM EGF and inversed the normalization by setting 0% as 100 and 100% as 0. These values were used to calculate the % of slowing down the rate of internalization.

To determine whether Lumi4Tb was not photo-bleached, we labelled SNAP-EGFR with Lumi4Tb and added tag-lite buffer instead of FB to the wells. We measured the 620 nm signal each 3 minutes for 120 minutes. The emission levels at 620 nm for Lumi4-Tb remained stable over time.

**Fluorescence microscopy**

Cells were prepared 48 hours before the start of the experiment. Lab-Tek®II 8-chamber #1.5 German Coverglass System plates (cat. no. 155409, Nunc) were coated with PLO and 25,000 cells per well in DMEM supplemented with 10% FBS were added. Cells were incubated at 37 °C and 5% CO_2_. At 4 hours before the start of the experiment, 200 nM of BG-Red in ice-cold serum-free DMEM was added and cells were incubated at 4 °C for 90 minutes. Next, wells were washed 4 times with Tag-Lite®-buffer and 250 µL ice-cold Tag-Lite®-buffer containing 1 µM erlotinib, 1 µM osimertinib, 10 nM cetuximab or vehicle was added and incubated for 2 hours at 4 °C. A pre-heated LSM780 confocal microscope (Zeiss) at 37 °C with 63X Plan Apochromat 1.4 NA oil DIC (Zeiss) and X-Cite 120LED Boost High-Power LED illumination system (Excelitas Technologies) was used for image acquisition. SNAP-Red was excited by a helium-neon laser at 633 nm. Images were taken in brightfield and red acceptor emission spectrum, right before addition of 250 µL pre-heated EGF (final concentration of 10 nM) or vehicle and after 30 minutes incubation at 37 °C. Image brightness was optimized by ImageJ (Version 2.1.0/1.53c, Fiji) for each condition. The same brightness settings were used for the same condition at 0 and 30 minutes.

**Data analysis**

All experiments have been performed at least three times and all graphs present the means and standard error of the mean (SEM) of at least three individually performed experiments unless stated otherwise. Curve fitting and statistical analysis was done with GraphPad Prism software (version 9.0) and is explained in the corresponding figure legends.

1.	P. Scholler, *et al.*, HTS-compatible FRET-based conformational sensors clarify membrane receptor activation. *Nat. Chem. Biol.* **13**, 372–380 (2017).

2.	A. Levoye, *et al.*, A broad G protein-coupled receptor internalization assay that combines SNAP-tag labeling, diffusion-enhanced resonance energy transfer, and a highly emissive terbium cryptate. *Front. Endocrinol. (Lausanne).* **6** (2015).
