## supplementary figures for "Different conformations of EGF-induced receptor dimers involved in signaling and internalization"

| A | EGFR per cell (relative to A431) | B | Labelling of SNAP-tag |
| --- | --- | --- | --- |
| 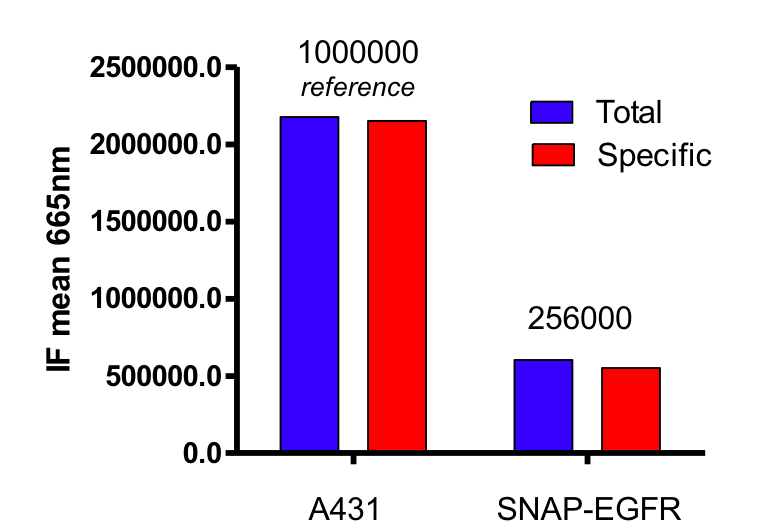 | | 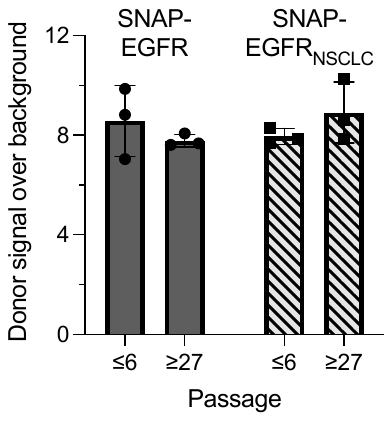 | |

**Suppl. Fig. S1.** Expression levels of SNAP-EGFR. *(A)* FACS analysis of EGF-red labelled EGFR. *(B)* Signal over noise of Lumi4-Tb labelled SNAP-EGFR. Data in *B* is mean ± SD of triplicates from a representative experiment.

| A | EGF | B | TGF-α |
| --- | --- | --- | --- |
| 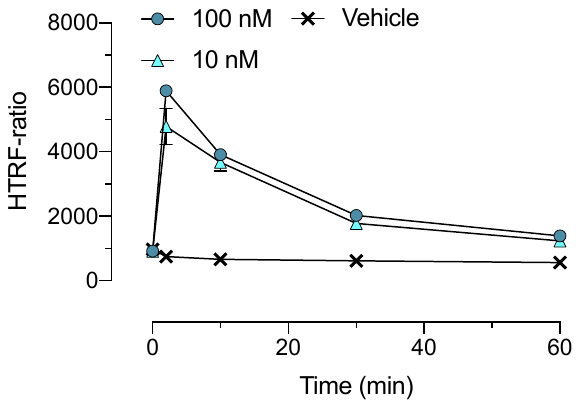 | | 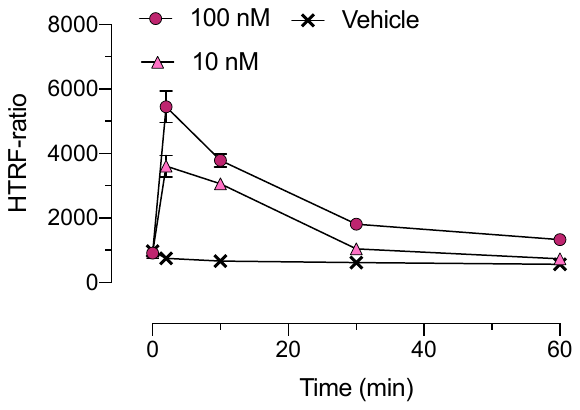 | |
| C | Dacomitinib | D | Erlotinib |
| 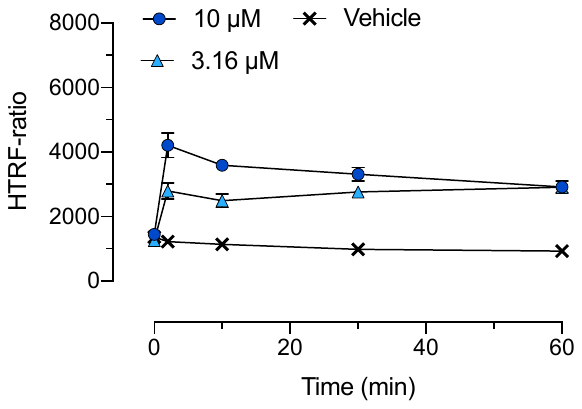 | | 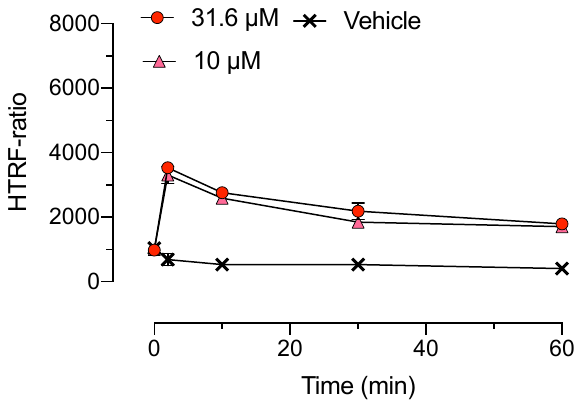 | |
| E | PD153035 | F | GW583340 |
| 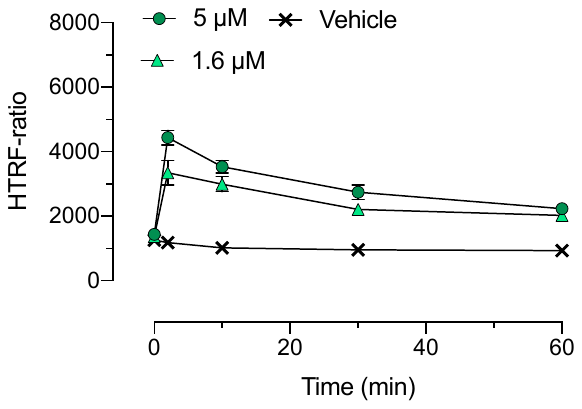 | | 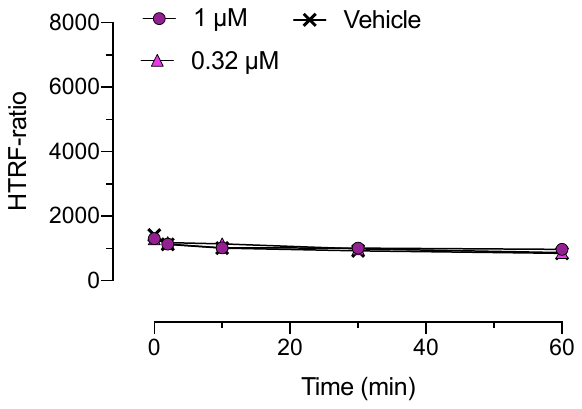 | |
| G | Lapatinib | H | Osimertinib |
| 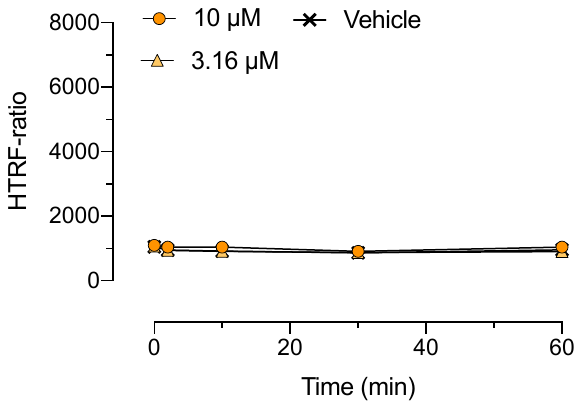 | | 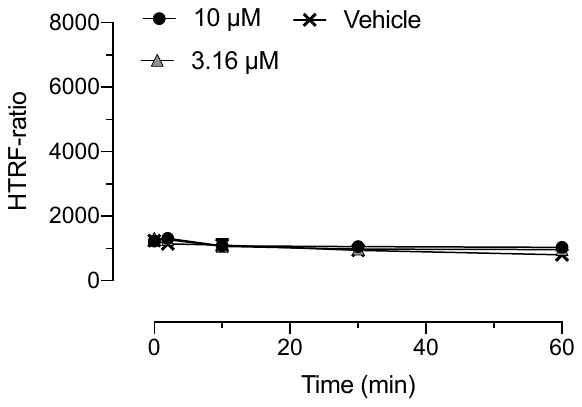 | |

**Suppl. Fig. S2.**	Intersubunit FRET for *wild-type* SNAP-EGFR. Intersubunit FRET is represented by HTRF-ratio. Data in *A-H* is mean ± SD of representative experiments.

| A | *wild-type* SNAP-EGFR  Intersubunit FRET | B | *wild-type* SNAP-EGFR (transient expression)  Intersubunit FRET |
| --- | --- | --- | --- |
| 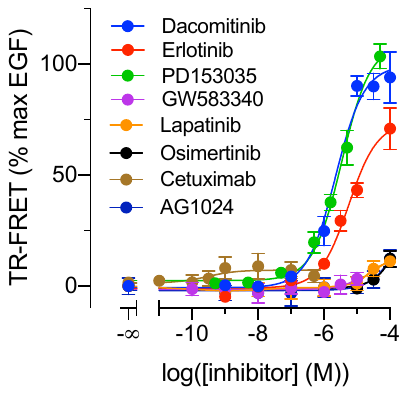 | | 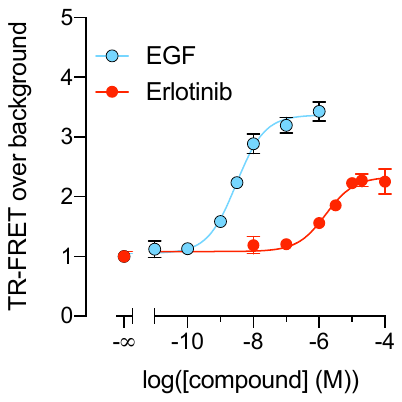 | |
| C | Dimerization EGFR_del242-249_  (transient expression)  Intersubunit FRET | D | SNAP-EGFR_delCter_  (transient expression)  Intersubunit FRET |
| 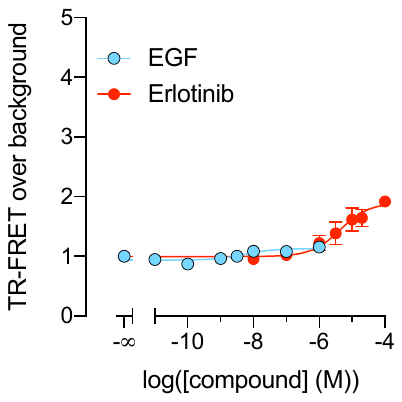 | | 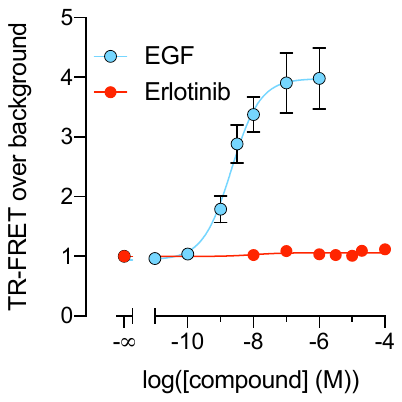 | |

**Suppl. Fig. S3.**	Intersubunit FRET assays. *(A)* TK inhibitor-induced intersubunit FRET. *(B)* Intersubunit FRET of transiently expressed *wild-type* SNAP-EGFR. *(C)* Intersubunit FRET of transiently expressed SNAP-EGFR_del242-249_. *(D)* Intersubunit FRET of transiently expressed SNAP-EGFR_delCter_. Data in *A-D* are mean ± SEM of at least three individual experiments.

| A | SNAP-mGlu_4_-C2 + CLIP-mGlu_2_-C1 | | |
| --- | --- | --- | --- |
| 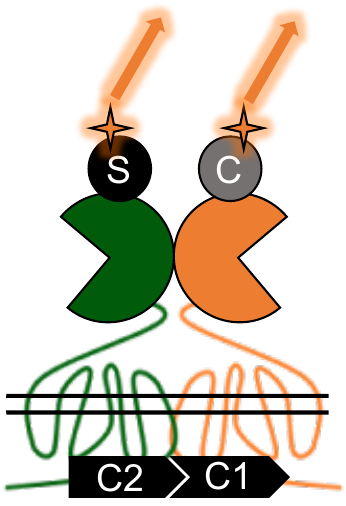 | | | 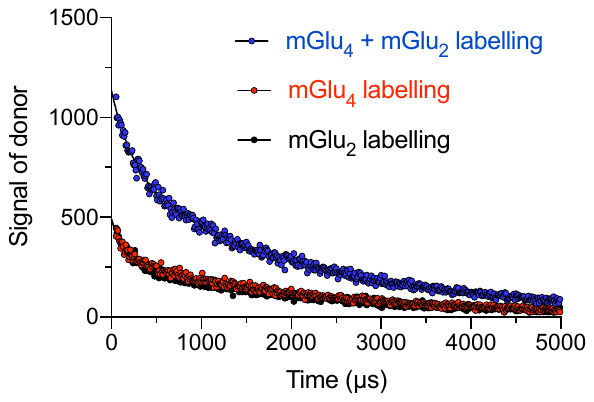 |
| B | | SNAP-EGFR | |
| 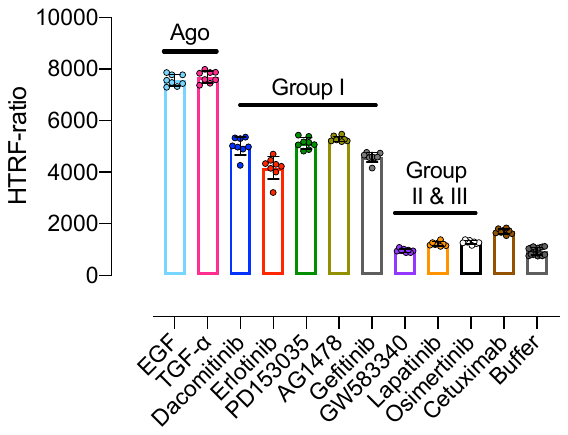 | | | |

**Suppl. Fig. S4.**	Expression levels of mGlu subunits in mGlu2-4 heterodimer and intersubunit FRET of *wild-type* SNAP-EGFR at 4 °C. *(A)* Scheme of mGlu2-4 labelling with donor and signal of the donor. *(B)* Intersubunit FRET of *wild-type* SNAP-EGFR at 4 °C represented by HTRF-ratio. Data in *A* are individual data points of a representative experiment. Data in *B* are mean ± SD of a representative experiment.

| A | Y1068 phospho | B | Y1068 phospho |
| --- | --- | --- | --- |
| 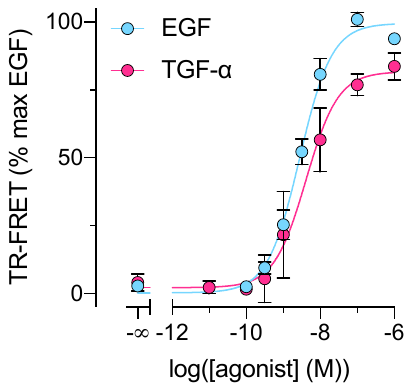 | | 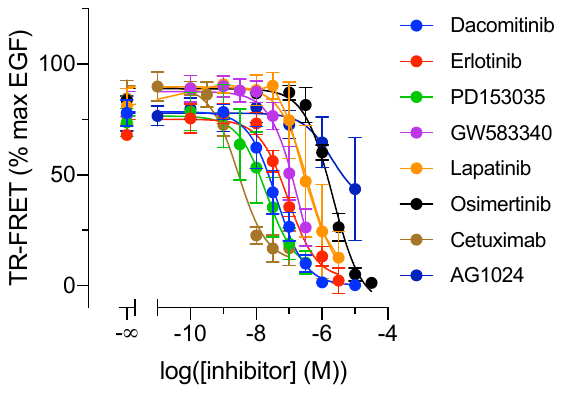 | |
| C | ERK1/2 phospho | D | ERK1/2 phospho |
| 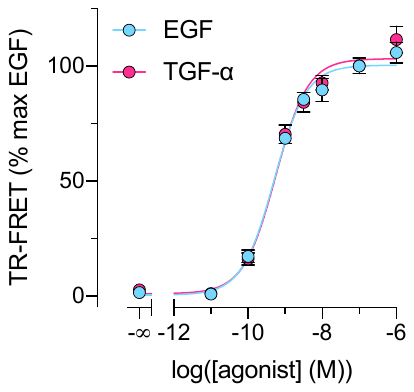 | | 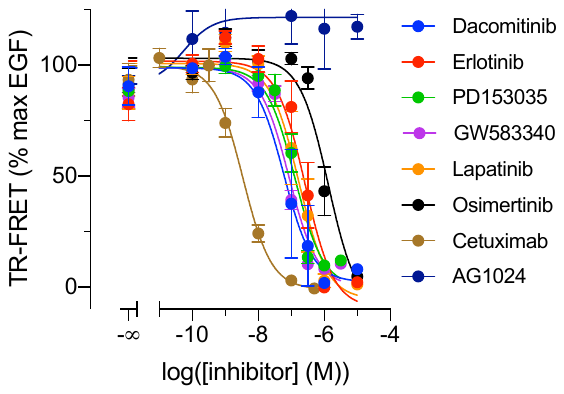 | |

**Suppl. Fig. S5.** Phosphorylation of Y1068 of the EGFR and Thr202/Tyr204 of ERK1/2. *(A)* Phosphorylation of Y1068 of the EGFR induced by agonists. *(B)* EGF-induced Y1068 of the EGFR phosphorylation (10 nM) inhibited by inhibitors. *(C)* Phosphorylation of ERK1/2 induced by EGFR agonists. *(D)* EGF-induced phosphorylation (10 nM) of ERK1/2 inhibited by inhibitors. Data in *A-D* are the mean ± SEM of at least three individual experiments.

+ osimertinib

| A | EGF  0 min | | B | Vehicle  0 min  30 min |
| --- | --- | --- | --- | --- |
| 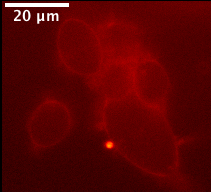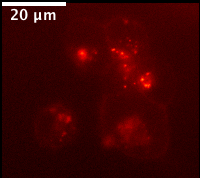  EGF  30 min  EGF | | | 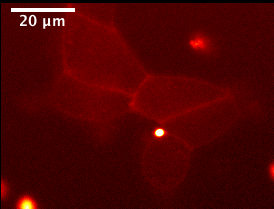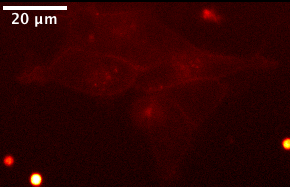  Vehicle  Vehicle | |
| C | EGF + erlotinib  30 min | | D | EGF + cetuximab |
| 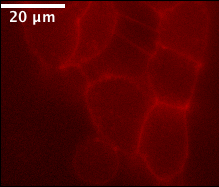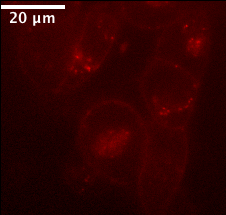  + erlotinib  + erlotinib  0 min | | | 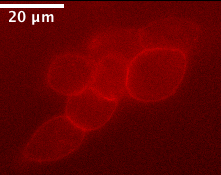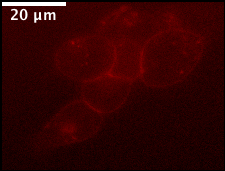  + cetuximab  + cetuximab  0 min  30 min | |
| E | | EGF + osimertinib | | |
| 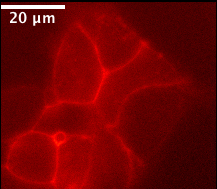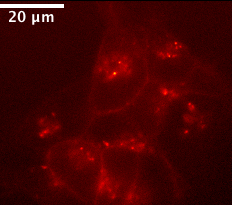  0 min  + osimertinib  30 min | | | | |

**Suppl. Fig. S6.** Internalization of the *wild-type* SNAP-EGFR labelled with SNAP-Red represented by two snapshots right before addition of compounds and 30 minutes after. *(A)* 10 nM EGF. *(B)* Vehicle. *(C)* 1 µM erlotinib pre-incubated for 2 hours and 10 nM EGF. *(D)* 100 nM cetuximab pre-incubated for 2 hours and 10 nM EGF. *(E)* 1 µM osimertinib pre-incubated for 2 hours and 10 nM EGF. Snapshots in *A-E* are from a representative experiment.

| A | EGF | B | TGF-α | C | EGF & TGF-α |
| --- | --- | --- | --- | --- | --- |
| 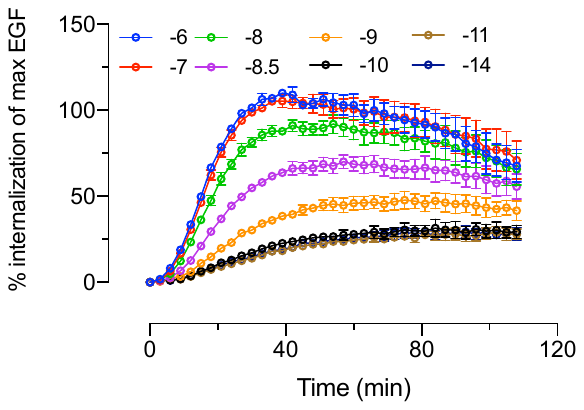 | | 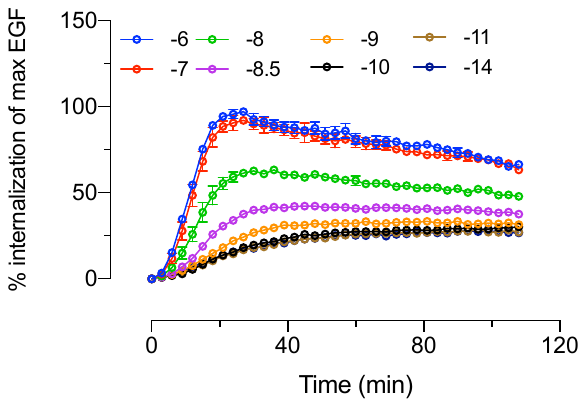 | | 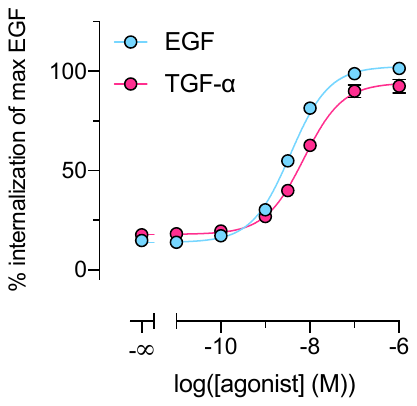 | |
| D | Dacomitinib | E | Erlotinib | F | PD153035 |
| 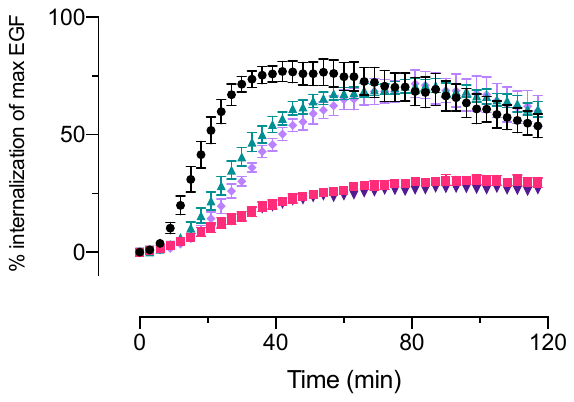 | | 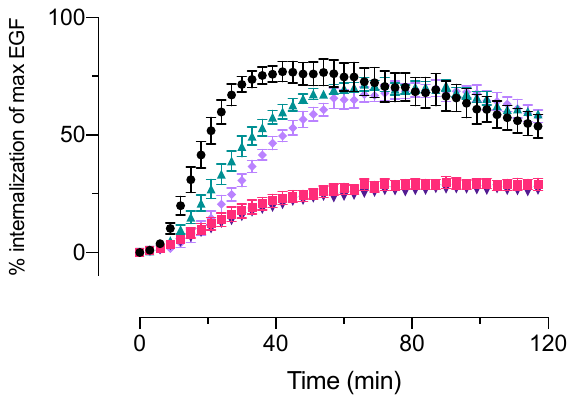 | | 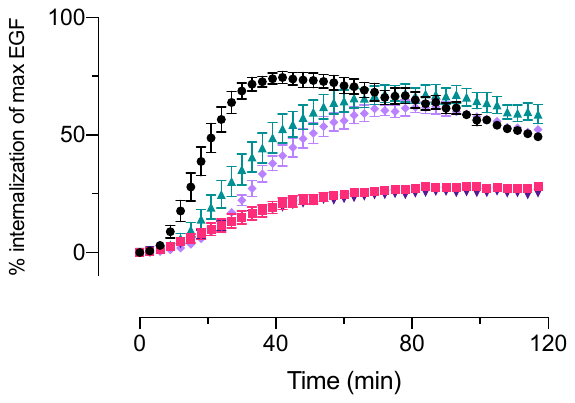 | |
| G | GW583340 | H | Lapatinib | I | Osimertinib |
| 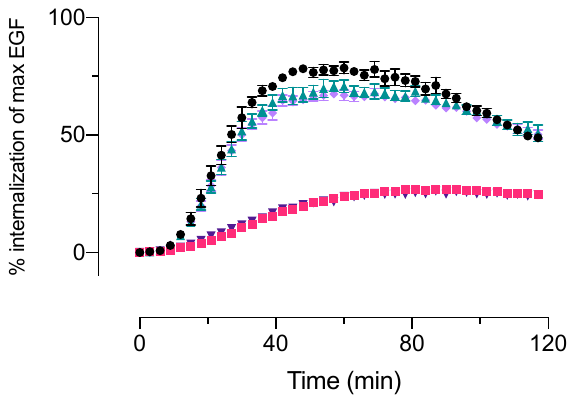 | | 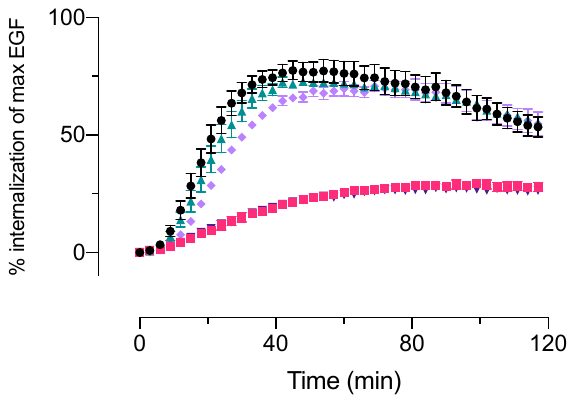 | | 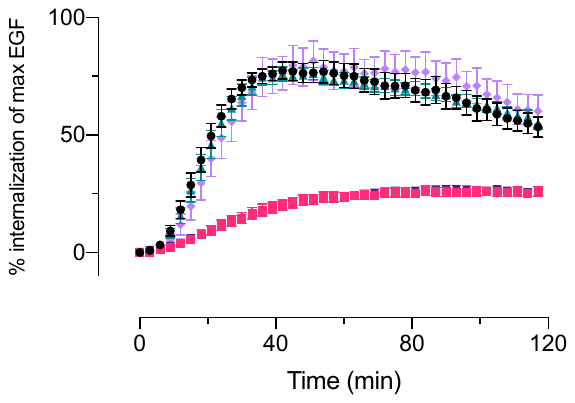 | |
| J | Cetuximab | K | AG1024 | L | Legend D-L |
| 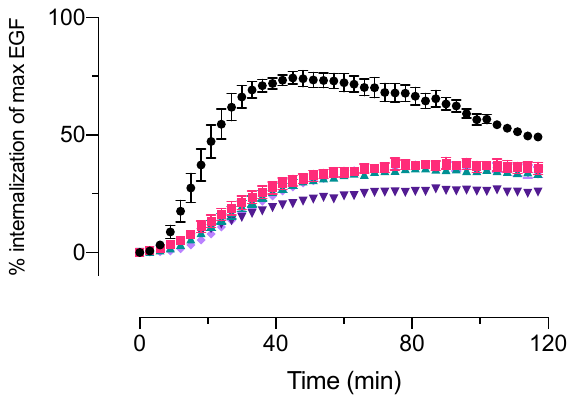 | | 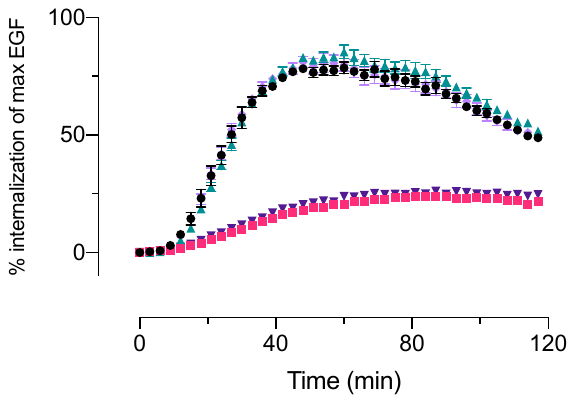 | | 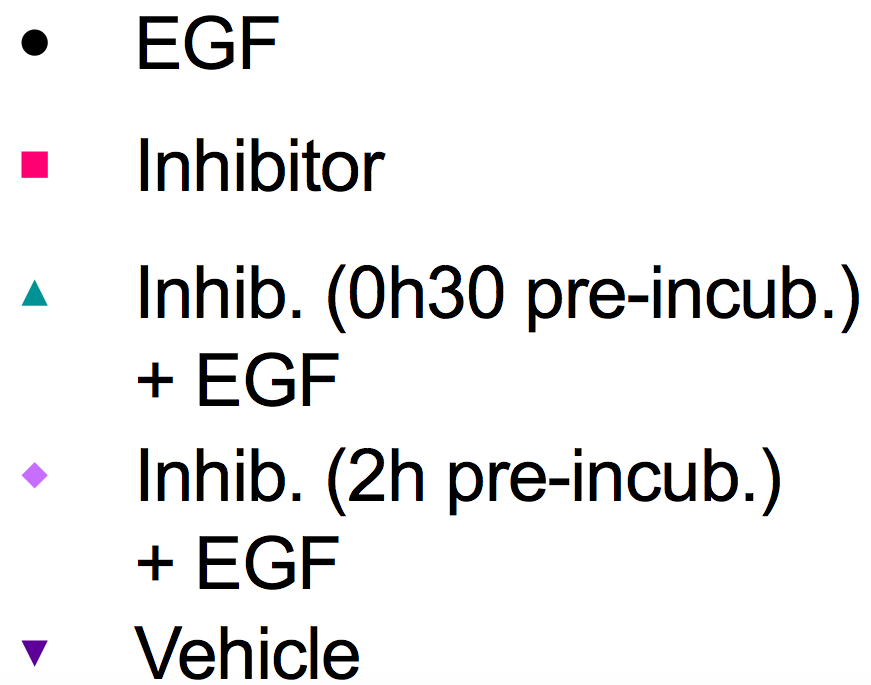 | |

**Suppl. Fig. S7.** Internalization of the *wild-type­* SNAP-EGFR labelled with SNAP-Lumi4-Tb represented by kinetic curves. *(A, B)* Agonists. *(C)* Dose-response curve of agonists. *(D-L)* Internalization of EGFR in presence of pre-incubated inhibitors and EGF. Data in A-B and D-K are mean ± SEM of at least three individual experiments.

| A | EGF | B | TGF-α |
| --- | --- | --- | --- |
| 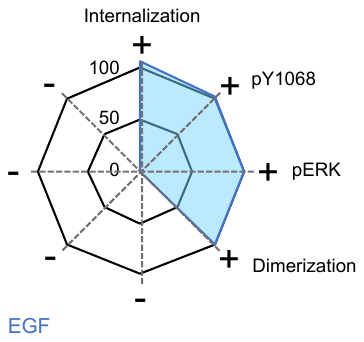 | | 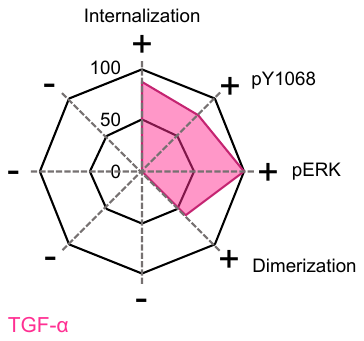 | |

**Suppl. Fig. S8.**	Pharmacological profile of compounds. (A) EGF. (B) TGF-α. All data are from figures 1-4.

| A | Int. FRET v. pY1068 | B | Int. FRET v. Internalization | C | Int. FRET v. pERK |
| --- | --- | --- | --- | --- | --- |
| 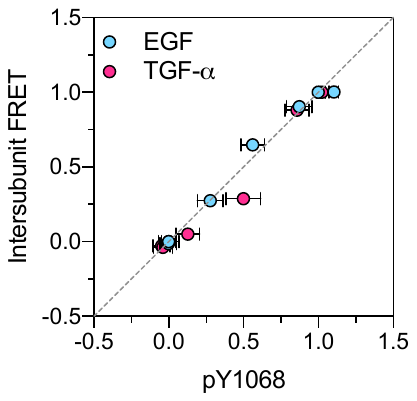 | | 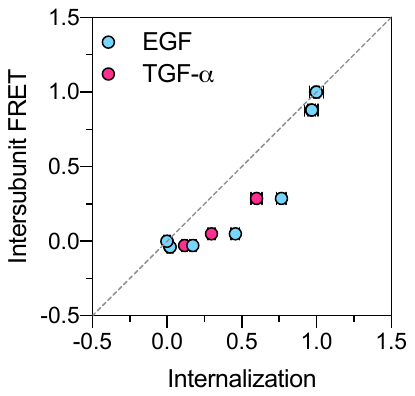 | | 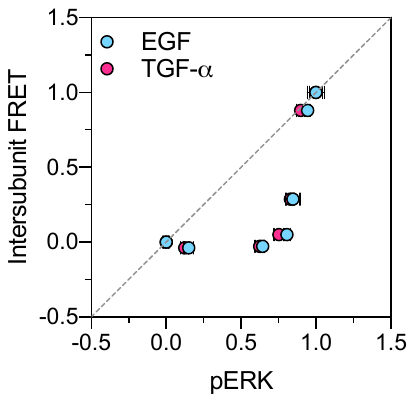 | |
| D | pY1068 v. pERK | E | pY1068 v. Internalization | F | Internalization v. pERK |
| 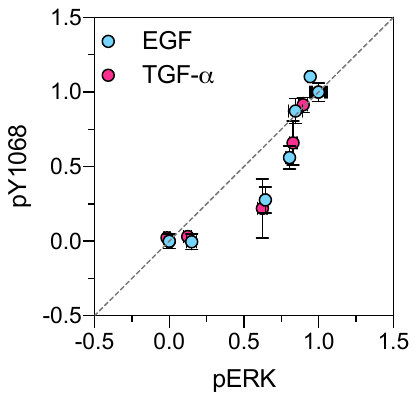 | | 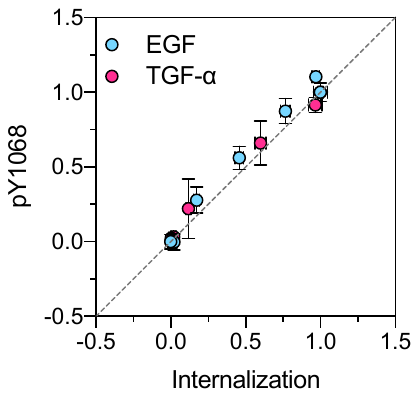 | | 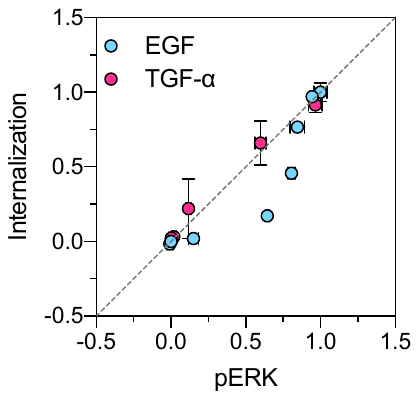 | |

**Suppl. Fig. S9.** Bias plots for EGFR agonists. *(A)* Intersubunit FRET v. phosphorylation Y1068 of the EGFR. *(B)* Intersubunit FRET v. Internalization. *(C)* Intersubunit FRET v. phosphorylation ERK. *(D)* phosphorylation Y1068 of the EGFR v. phosphorylation of ERK1/2. *(E)* phosphorylation Y1068 of the EGFR v. Internalization. *(F)* Internalization v. phosphorylation Y1068 of ERK1/2. All data are mean ± SEM of at least three individual experiments.

| A | Dacomitinib | B | Erlotinib | C | PD153035 | D | GW583340 |
| --- | --- | --- | --- | --- | --- | --- | --- |
| 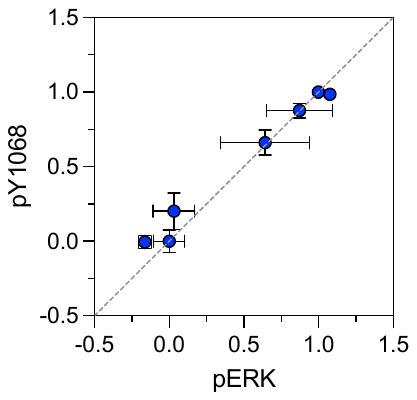 | | 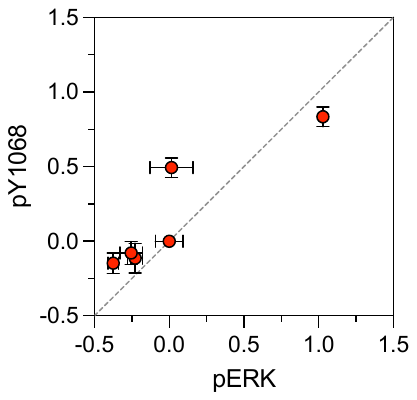 | | 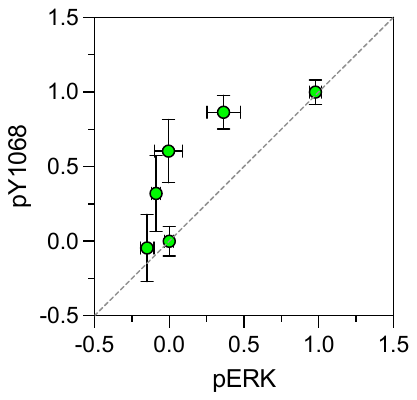 | | 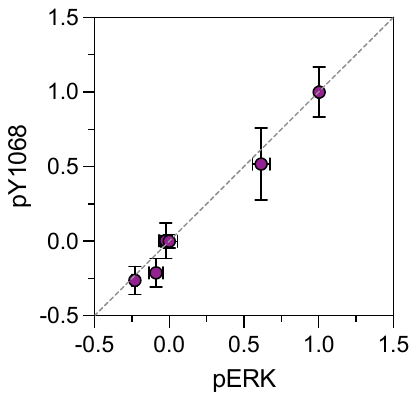 | |
| E | Lapatinib | F | Osimertinib | G | Cetuximab |  | |
| 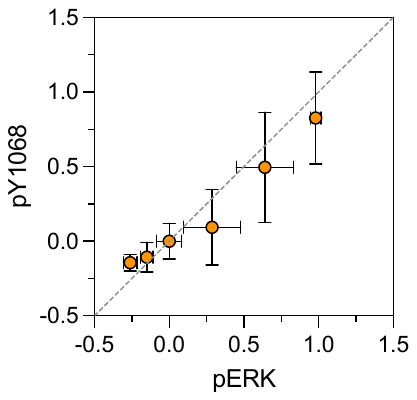 | | 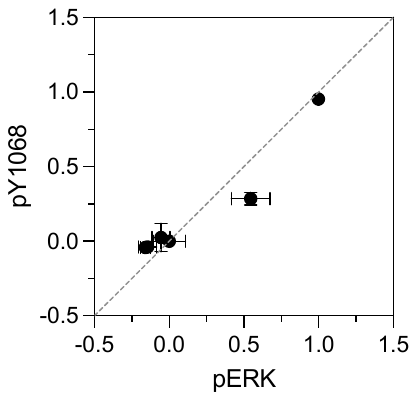 | | 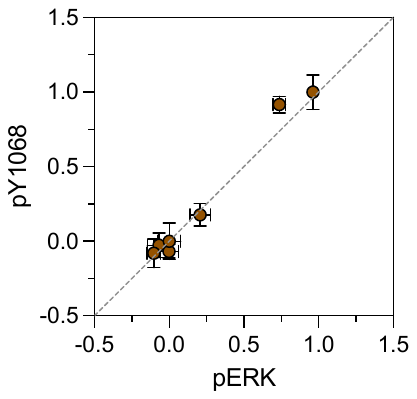 | |  | |

**Suppl. Fig. S10**. Inhibition of EGF-induced phosphorylation of the EGFR or ERK1/2 by TK inhibitors.

| A | WT(*CLIP*)-NSCLC(*SNAP*) | B | NSCLC(*SNAP*) |
| --- | --- | --- | --- |
| 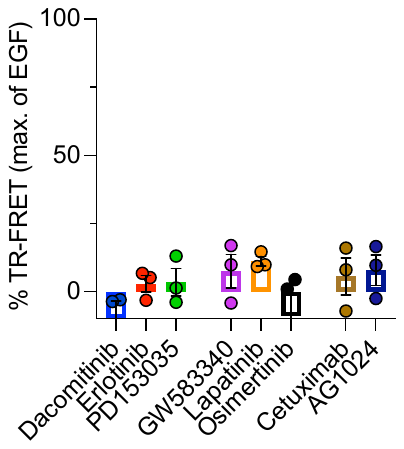 | | 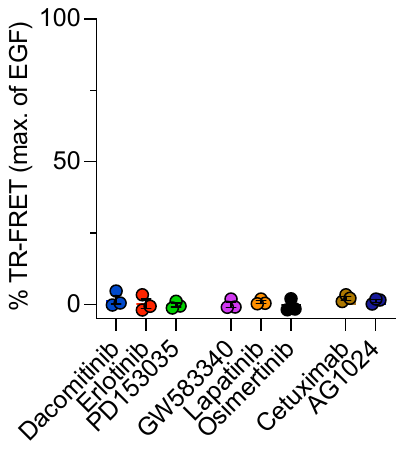 | |

**Suppl. Fig. S11.** Effect of inhibitors on *wild-type* EGFR – EGFR_NSCLC_ intersubunit FRET assay. *(A)* *Wild-type* CLIP-EGFR + SNAP-EGFR_NSCLC_ heterodimer. *(B)* SNAP-EGFR_NSCLC_ homodimer. Data A-B are mean ± SEM of three individual experiments.

**Supplementary tables**

| Ligand | α_DA_ | SD | A_s_ | SD |
| --- | --- | --- | --- | --- |
| AG1478 | 0.84 | 0.04 | 0.40 | 0.07 |
| Dacomitinib | 0.85 | 0.06 | 0.49 | 0.11 |
| Erlotinib | 0.84 | 0.06 | 0.41 | 0.08 |
| Gefitinib | 0.84 | 0.05 | 0.43 | 0.10 |
| PD153035 | 0.84 | 0.03 | 0.38 | 0.08 |
| GW583340 | 0.82 | 0.03 | 0.16 | 0.02 |
| Lapatinib | 0.75 | 0.09 | 0.16 | 0.06 |
| Osimertinib | 0.81 | 0.12 | 0.25 | 0.13 |
| EGF | 0.75 | 0.04 | 0.31 | 0.07 |
| TGF-α | 0.77 | 0.04 | 0.35 | 0.04 |
| Cetuximab | 0.88 | 0.04 | 0.30 | 0.11 |
| Vehicle | 0.80 | 0.15 | 0.20 | 0.11 |

**Suppl. Table S1.** The true fraction of the slow decay species (α_DA_) after the Heyduk and Heyduk correction.

| pEC50 ± SEM | EGF | N | TGF-α | N |
| --- | --- | --- | --- | --- |
| EGFR_WT_ | 8.636 ± 0.039 | 9 | 7.636 ± 0.019 | 3 |
| EGFR_NSCLC_ | 8.705 ± 0.204 | 4 | 8.131 ± 0.164 | 3 |
| WT + NSCLC | 8.384 ± 0.040 | 3 | 7.484 ± 0.251 | 3 |
| P value | NS (P = 0.1690) |  | NS (P = 0.0860) |  |

**Suppl. Table S2.** Summary of potencies of EGF and TGF-α for EGFR_WT_, EGFR_NSCLC_ and EGFR_WT_ + EGFR_NSCLC_. Potency values are represented as pEC50 ± standard error of the mean (SEM) of 3 or more individual experiments. The used statistical test is ordinary one-way ANOVA. NS means not significant.

| **Ligand** | **Mean ± SD** | **N** | **P value** |
| --- | --- | --- | --- |
| Erlotinib | 405.2 ± 57.0 | 24 | 0.6972 |
| Gefitinib | 412.9 ± 52.8 | 24 | 0.8606 |
| PD153035 | 383.1 ± 33.8 | 24 | 0.2186 |
| AG1478 | 384.6 ± 43.9 | 24 | 0.2408 |
| Dacomitinib | 414.3 ± 48.3 | 24 | 0.8844 |
| GW583340 | 828.2 ± 133.8 | 24 | <0.0001 |
| Osimertinib | 601.6 ± 199.2 | 23 | 0.0002 |
| Lapatinib | 549.2 ± 142.2 | 24 | 0.0483 |
| EGF | 456.8 ± 72.8 | 84 | - |
| TGF-α | 382.0 ± 46.6 | 24 | 0.2016 |
| Cetuximab | 733.3 ± 167.6 | 24 | <0.0001 |
| Vehicle | 812.3 ± 248.4 | 84 | <0.0001 |

**Suppl. Table S3.** Summary of basal, TK inhibitor-induced, agonist-induced mean excited-state lifetimes of sensitized acceptor emission. Statistical test is one-way ANOVA with Dunnett’s multiple comparison test.
